# Biodiversity forecasting in natural plankton communities reveals temperature and biotic interactions as key predictors

**DOI:** 10.1101/2024.10.10.617676

**Authors:** Ewa Merz, Francesco Pomati, Serguei Saavedra, Luis J. Gilarranz

**Affiliations:** Department of Aquatic Ecology, Eawag (Swiss Federal Institute of Aquatic Science and Technology) Überlandstrasse 133, 8600, Dübendorf, ZH, Switzerland; Scripps Institution of Oceanography, University of California San Diego, 92037 La Jolla, CA, USA; Department of Civil and Environmental Engineering, MIT, 77 Massachusetts Av, 02139 Cambridge, MA, USA; Santa Fe Institute, Santa Fe, NM 87501, USA

**Author notes:** These authors contributed equally (Francesco Pomati, Luis J. Gilarranz).

## Abstract

As natural ecosystems experience unprecedented human-made degradation, it is urgent to deliver quantitative anticipatory forecasts of biodiversity change and identify relevant biotic and abiotic predictors. Forecasting natural ecosystems has been challenging due to their complexity, chaotic nonlinear nature and the availability of adequate data. Here, we use four years of daily abundance of a complex lake planktonic ecosystem and its abiotic environment to model and forecast biodiversity metrics. Using a state-of-the-art equation-free modelling technique, we forecast community richness and turnover with a proficiency greater than the constant predictor several generations ahead (30 days). Short-term forecasts improve substantially using biotic predictors (i.e., autoregressive term or community richness). Long-term forecasts require a more complex set of variables (i.e., biotic interactions), and the forecast proficiency depends strongly on including abiotic predictors such as water temperature. Depending on the forecast horizon, biotic and abiotic predictors can interact nonlinearly and synergistically, enhancing each other’s effects on biodiversity metrics. Our findings showcase the challenges of forecasting biodiversity in natural ecosystems and stress the importance of monitoring focal biotic and abiotic predictors to anticipate undesired changes.

## Introduction

Accurate biodiversity forecasts for global and local disturbances are essential to effective decision-making and conservation strategies in a rapidly changing world^1,2^. This forecasting effort requires a balance between mechanistic models, which allow an explanation of ecological processes, and robust models that anticipate future change. While mechanistic models have been the foundation of ecological research, their complexity often hinders practical forecasting due to the high-dimensional, interdependent, and non-linear nature of ecological systems^3,4^. Anticipatory predictions based on the system’s history (i.e., forecasting), as opposed to explanatory predictions by mechanistic models, are promising for biodiversity forecasting given the above limitations and the increasing socio-political demand for actionable insights into global environmental changes^5,6^.

Two primary approaches guide biodiversity forecasting: species distribution models (SDMs) and macroecological models (MEMs)^1^. SDMs forecast future species’ spatial ranges or temporal trajectories based on ecological niche models built from current data and projected climate scenarios^7^. By summing individual species forecasts, they estimate species richness and composition across landscapes or over time^8^. However, SDMs typically focus on individual species abundances, struggle with rare species due to limited data, and often misestimate biodiversity metrics when stacking individual species predictions^1,8^. Additionally, species’ realised niches can be narrower than their potential environmental ranges due to biotic interactions, which breaks a core assumption of SDMs^9^. MEMs, in contrast, are statistical (e.g., regression, machine learning) models used to fit and forecast biodiversity metrics based on environmental variables. MEMs provide less computationally expensive and biased biodiversity forecasts compared to stacked SDMs^1,7^.

In both SDM and MEMs, biotic factors (e.g., interactions among species) can significantly improve biodiversity forecasts^10^. However, adding other species, simple biotic variables or species interactions as predictors in models represents a challenge in biodiversity forecasting due to the shortage of available data and the high multidimensionality of required models^5,11^. Only a few cases have accounted for biotic interactions explicitly^7,12–14^. Integrating abiotic drivers of communities, such as temperature and other climatic variables, alongside biotic factors, such as ecological interactions, remains a critical research gap for enhancing forecast accuracy^15,16^.

Biodiversity forecasting approaches also face other challenges. High goodness-of-fit in model calibration does not necessarily translate to accurate forecasts. This discrepancy highlights the necessity of long-term and high-resolution data for model calibration and robust validation^17^. In general, the intrinsic unpredictability of complex adaptive systems, characterised by chaotic dynamics (bounded, deterministic, aperiodic fluctuations that display sensitive dependence on initial conditions), which makes long-term (but not short-term) prediction impossible, challenges the reliability of long-term forecasts of biodiversity^18,19^. The influence of chaotic dynamics in biodiversity forecasts has been overlooked, perhaps due to the focus on single-species models. Chaos has been demonstrated under laboratory and natural conditions in many communities that are a target for conservation or play a crucial role in important ecosystem services including plankton, macroinvertebrates, insects, and mammals^20^. Chaotic dynamics are generally driven by fluctuations in both abiotic (e.g., temperature or nutrients) and biotic factors (e.g., biotic interactions)^20^, and how these interact to affect biodiversity forecast at different temporal scales remains untested.

Here, we use a MEM approach based on state space reconstruction (SSR) methods to forecast biodiversity metrics in a lake plankton community using abiotic and biotic factors, studying their relative importance and how they interact to affect forecasts at different horizons. SSR methods do not rely on detailed mechanistic knowledge of the focal system for forecasting and outperform traditional models in chaotic systems^4,21^, like plankton communities^19^. Despite their relevance for aquatic ecosystem processes and services, plankton are a notable worst-case scenario in terms of predictability: they have a high diversity of interacting taxa and fast generation time, they are sensitive to internally driven community dynamics (e.g. biotic interactions), stochastic perturbations, and they are forced by external seasonal changes in temperature and nutrients that modify the ecosystem’s behaviour over time^19,22–24^.

Here, we employ regularised S-maps to forecast plankton biodiversity metrics based on network properties and abiotic environmental conditions up to 30 days ahead (**Fig. 1**). S-maps are multilinear regressions weighted in state space, which allows for complex, nonlinear relationships (i.e., estimated coefficients) between variables to vary over time^25–27^. Regularisation techniques control for model overfitting and account for collinearity between explanatory variables^26^. We choose a 30-day forecast horizon to allow comparison with previous work forecasting plankton communities^24,28,29^, and it represents roughly 3 to 30 generations for the slowest (mesoplankton) and the fastest (nano-microplankton) growing organisms in the community, respectively.

**Fig. 1:**
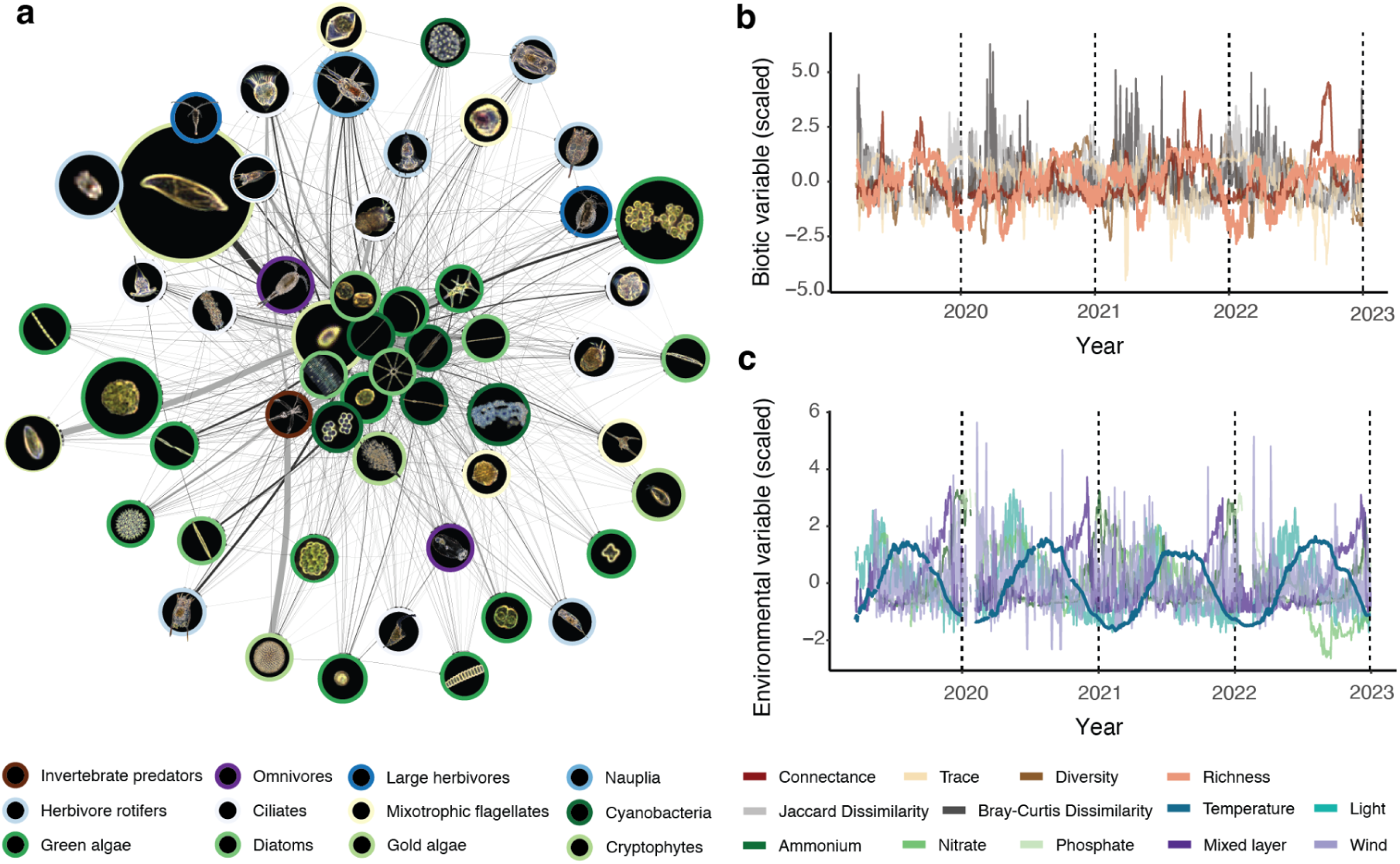
Forecasting biodiversity change in a lake plankton community. **[a]** Interaction network between all plankton taxa, reconstructed using multiview distance S-maps and 4 years of daily abundance data: the line thickness represents the magnitude of the effect of one taxon’s abundance on another (from the Jacobian interaction matrix), averaged over time. For the species list see **Table S2**. From this network, we derive connectance (the number of realised nodes) and the trace of the Jacobian matrix (which quantifies the sensitivity of a community to environmental fluctuations). We forecast change in community diversity metrics (e.g. richness and Jaccard dissimilarity in community composition) **[b]** as a function of biotic ecosystem properties (e.g. connectance, trace) **[b]** and **[c]** abiotic variables (water temperature, light penetration, concentrations of ammonium, nitrate, and phosphate, depth of the mixed layer, and maximum daily wind speed).

In this study, we tracked daily taxa abundances for the entire planktonic ecosystem of Lake Greifen (Switzerland) for four consecutive years (1332 days). We used an automated underwater microscope (**Methods**) to record all organisms from 10 μm to 1 cm, from invertebrate predators, large and small herbivores, and mixotrophic to photosynthetic protists, for a total of 55 taxa. A neural network classified every individual object detected to obtain the time series of taxa abundances at the daily temporal resolution (**Tab. S1, Fig. S1**). At the same location, we collected physical and chemical parameters that influence changes in plankton community abundance and composition^23^, i.e., water temperature, resources, and indicators of turbulent mixing (**Fig. 1, Fig. S2**).

We used multiview-distance regularised S-maps to infer time-varying biotic interactions in the plankton communities (**Methods**, **Fig. 1**). The multiview-distance add-on allows the S-map to accommodate many explanatory variables^27^. The estimated coefficients of the S-map relate to the Jacobian interaction matrix, indicating how a change in one taxon’s abundance will result in a change in another taxon’s abundance. Using the S-map’s coefficients, we calculated network connectance (the percentage of realised links at a given time) and the trace of the Jacobian matrix (a measurement of the community’s sensitivity to environmental perturbations, based on the strength of interactions)(**Fig. S3**)^27,30^. Properties of the interaction network can signal large-scale changes in the whole ecosystem^31^. Here, we include connectance and trace of the Jacobian since the former is a common indicator of the number while the latter summarises the strength of biotic interactions^32,33^.

We used the collected biotic and abiotic variables to forecast plankton biodiversity, which was estimated using four common metrics. The metrics are of two types, based on i) presence-absence data (taxonomic richness and Jaccard dissimilarity, **Fig. 2**), and ii) taxa relative abundances (Shannon diversity and Bray-Curtis dissimilarity, **Fig. E1**). Testing for chaos in time series using Lyapunov exponents, we find that dissimilarity metrics show evidence for chaotic dynamics, while richness metrics do not (**Fig. E2**). Here, we show the results of richness and Jaccard dissimilarity, and report results of abundance-based metrics in the **Extended Data** because the former, being based on presence-absence data, are the most commonly used metrics of biodiversity and compositional turnover^34^ and because, as we shall see, they are the most predictable. Note that we used biotic and abiotic predictors at a one-day lag relative to the response variable, even for the longest-term forecasts: the goal was to capture contemporary relationships and study how they allow forecasting at different horizons.

**Fig. 2:**
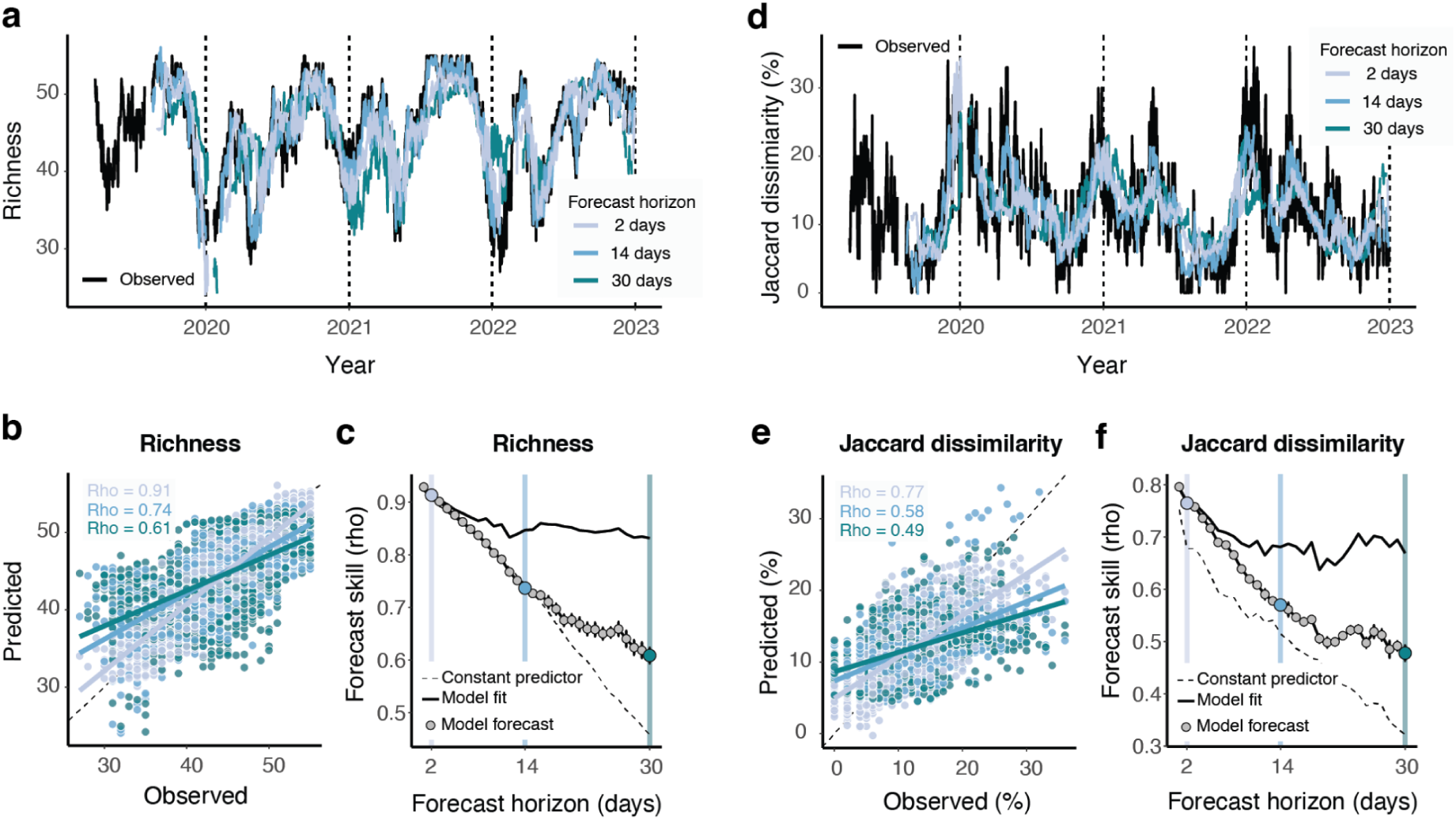
Forecast proficiency when predicting biodiversity change (richness and turnover) based on biotic and abiotic factors. **[a]** Time series of observed and predicted values for the forecast horizons 2, 14 and 30 days. **[b]** Correlation between predicted and observed values: the dotted line indicates the 1 to 1 line. **[c]** Forecast skill (estimated as the Pearson correlation coefficient between observations and predictions) of the six best models (average ± standard deviation) decreases with the forecast horizon: in the case of Richness (non-chaotic dynamics [**Fig. E2**]), the model outcompetes the constant predictor only after 14 days ahead forecasts; in the case of Jaccard dissimilarity (chaotic dynamics), the model always performs better than the constant predictor. The model fit shows the model’s performance used to study the interactions in Fig. 4.

## Results and discussion

### Forecasting proficiency of four common biodiversity metrics

Forecasting models of biodiversity and compositional turnover outcompete the constant predictor (baseline model that predicts constant diversity) across all tested temporal scales, spanning 1 to 30 days ahead (**Fig. 2**). The model uncertainty (standard deviation based on averaging the six best models - **Methods**), is very low. Outperforming the constant predictor several generations ahead shows that the data incorporated information on the processes that link biodiversity change to biotic and abiotic environmental factors. On the other hand, our approach shows limitations for abundance-based biodiversity metrics like Shannon diversity and Bray-Curtis dissimilarity (**Fig. E1**). Measurement errors and stochastic fluctuations in abundances, particularly prevalent in rare taxa, amplify when combined into community metrics, making abundance-based metrics challenging to model. More accurate estimates of abundances for rare taxa, filtering or accounting for measurement noise^35^, or smoothing of high-frequency plankton time series are required to improve the forecast proficiency relative abundance-based biodiversity metrics.

Our presence-absence-based models’ proficiency in forecasting plankton biodiversity change is comparable to or higher than previous work based on experimental communities^24^ and observational data^36^ targeting the abundances of aggregated taxonomic units. For example, only a few plankton functional groups, which are supposed to show high predictability^37^, have displayed forecast skills > 0.5 for a 30-day horizon. Our models outperform previous forecasting studies based on bulk phytoplankton community measures, which reached skills of 0.5 after 7 days^38^. Our models also deliver a better proficiency than former MEMs in forecasting richness in other systems^11,39,40^. We note that most previous studies use hindcasting, where models recreate past conditions, rather than actual forecasting, with rare exceptions^11^. While forecast proficiency decreases with the forecast horizon as expected^2,24,28^, we observe two novel patterns: i) the dissimilarity indices forecasts skill (chaotic time series) depart from the constant predictor all the time, while the richness forecast skill (non-chaotic) outcompete the constant predictor only after 14 days, and ii) the forecast skill’s ’ distance to the constant predictor increases with the forecast horizon (**Fig. 2c, f, Fig. E3**). These patterns suggest that richness time series might suffice relatively simple models (e.g., autoregressive) for short-term forecasting; dissimilarity time series may require more complex models. Biotic and abiotic predictors become more important for the forecasting models’ performance with increasing forecast horizons.

### Importance of biotic and abiotic predictors

We find that forecast proficiency relies mostly on a small set of biotic and abiotic predictors, whose importance (measured as the decrease in forecast skill when we omit the focal variable from the forecast model) changes as a function of the forecast horizon (**Fig. 3**). Short-term forecasting of plankton biodiversity relies on biotic factors (autoregressive term or community properties like richness)^41^. Long-term forecasts require a more complex set of variables (including indicators of biotic interactions), and the forecast proficiency crucially depends on the trend of the most important abiotic factor - temperature. For example, in long-term forecasting (30 days in our study) of community compositional turnover, temperature and biotic interactions (network connectance) are more important than baseline richness levels (**Fig. 3b**).

**Fig. 3:**
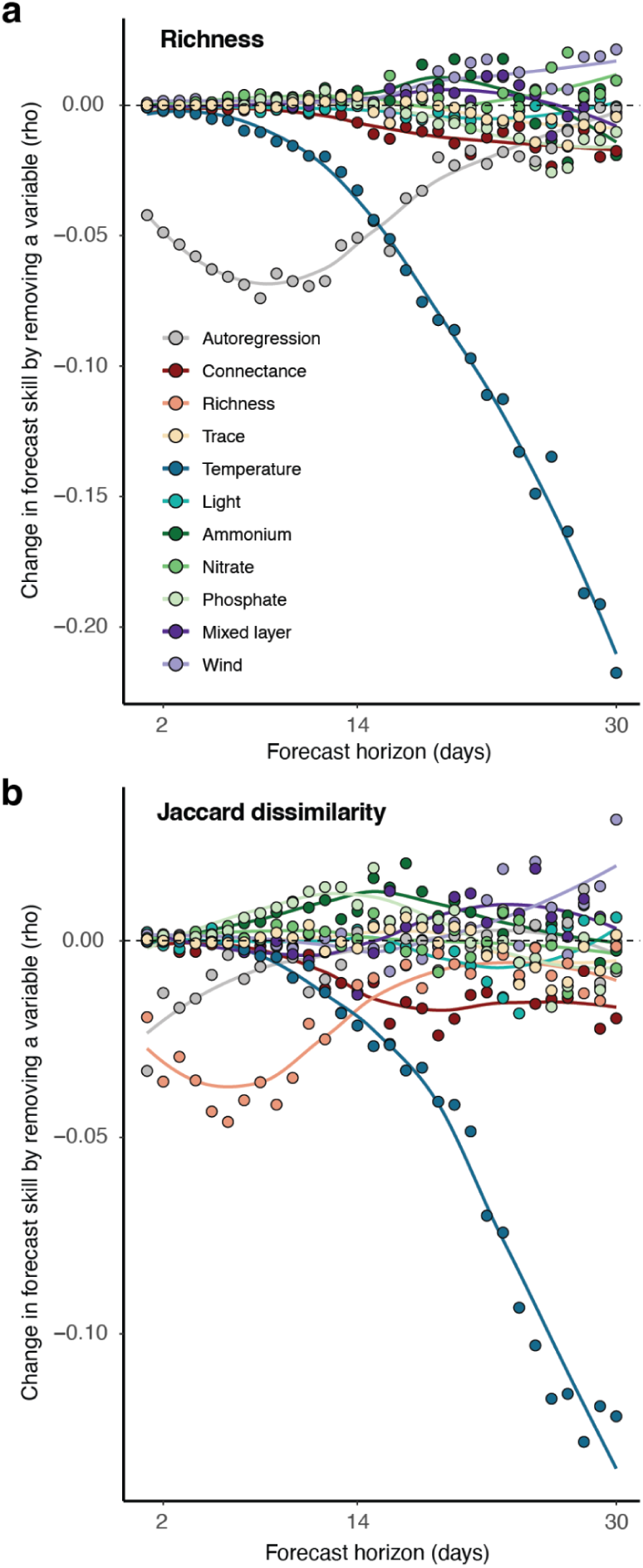
The magnitude and relative importance of biodiversity predictors change as a function of the forecast horizon. **[a]** Richness; **[b]** Jaccard dissimilarity. The y-axis shows the change in forecast skill (rho, Pearson’s correlation) when a variable is removed from the S-map’s model for each forecast horizon. The splines are a visual aid to show the trends in change in forecast skill as a function of the forecast horizon. Note that negative values indicate the magnitude of importance of each variable for forecast proficiency, while positive values mean that the model gains skill when the variable is omitted.

Water temperature is the most important variable for mid to long-term forecasting of all biodiversity metrics (**Fig. E4, Fig. E5**). Water temperature directly affects plankton physiology, ecology, and phenology^42,43^, and indirectly affects water physics (stability and turbulence of the water column) and chemistry (nutrient upwelling)^44,45^. In general, temperature is a crucial predictor of biodiversity change across systems and spatiotemporal scales (e.g. seasonal succession and climate change)^23,46–48^. It also acts in concert with additional abiotic factors and biotic interactions, which are rarely studied^49^. Resource levels play an important role in long-term forecasts of Shannon diversity and Bray-Curtis dissimilarity in our plankton communities, indicating that nutrients may be important when forecasting abundance-based but not presence-absence metrics of biodiversity (**Fig. E4, Fig. E5**). Most of the other abiotic variables did not affect the forecast proficiency of plankton biodiversity models, despite previous knowledge and their general use for modelling plankton dynamics^50^. For example, meteorological conditions (e.g. wind speed) do not improve forecasts and, indeed, might worsen forecast proficiency (e.g., by introducing noise) for both taxonomic richness and Jaccard dissimilarity (**Fig. 3**).

The connectance of the interaction network, and indicator of the number of biotic interactions, is an important factor for long-term forecasts of plankton biodiversity indices (**Fig. 3**, **Fig. E4, Fig. E5**). Our results confirm the inclusion of indicators of biotic interactions improve biodiversity forecast proficiency, as previously advocated^13–16^, specifically for presence-absence-based metrics (**Fig. 3**). By considering biotic interactions and how they change over time, our modelling approach acknowledges that biodiversity is an emergent property of a complex system of interacting agents and shows how this information is necessary to anticipate its future states. Chaotic time series (e.g., Jaccard dissimilarity) depend more on such internal factors (like connectance, which emerges from non-linear interactions and ecological feedback) compared to non-chaotic time series (richness) (**Fig. 3**, **Fig. E4, Fig. E5**). However, biotic interactions and emergent network properties can depend on abiotic environmental conditions^31^.

### Interactions between biotic and abiotic predictors

Finally, we aim to explore the nature (linear versus nonlinear) and effect (strength and direction) of interactions between abiotic and biotic predictors on biodiversity metrics at various forecast horizons. Such information is crucial to understanding and improving the proficiency of biodiversity forecasting models. We manipulate the biodiversity indices’ strongest abiotic and biotic predictor (from **Fig. 3**) from low to high values while keeping other variables constant to archive this (**Methods**). Based on those values, we forecast biodiversity metrics (richness and Jaccard dissimilarity) 1 to 30 days ahead.

The strength, direction, and nonlinearity of the emergent interactions between biotic and abiotic predictors change with the forecast horizon (**Fig. 4**). At short forecast horizons, abiotic predictors do not affect biodiversity indices, or we find no interaction between biotic and abiotic factors (**Fig. 4 a, e, h**). For example, we predict higher Jaccard dissimilarity when taxonomic richness is high, independent of water temperature (**Fig. 4 e, h**). At medium forecast horizons, biotic and abiotic predictors display synergistic nonlinear interactions, enforcing each other’s effect on biodiversity metrics (**Fig. 4 b, f, d, h**). Richness is predicted to be highest at high water temperature and network connectance, while Jaccard dissimilarity is lowest at high water temperature and taxonomic richness (**Fig. 4 d, f)**. These results support prior evidence from lakes showing that, with increasing water temperature (climate warming), plankton communities show higher richness^46^ and lower turnover^51,52^. Besides plankton communities, increasing temperature has been associated with increased richness^53–56^ and decreased turnover rates in other ecosystems^57,58^. At long-term forecast horizons, interactions between biotic and abiotic predictors are weak for both biodiversity metrics (**Fig. 4 c,d,g,h**).

**Fig. 4:**
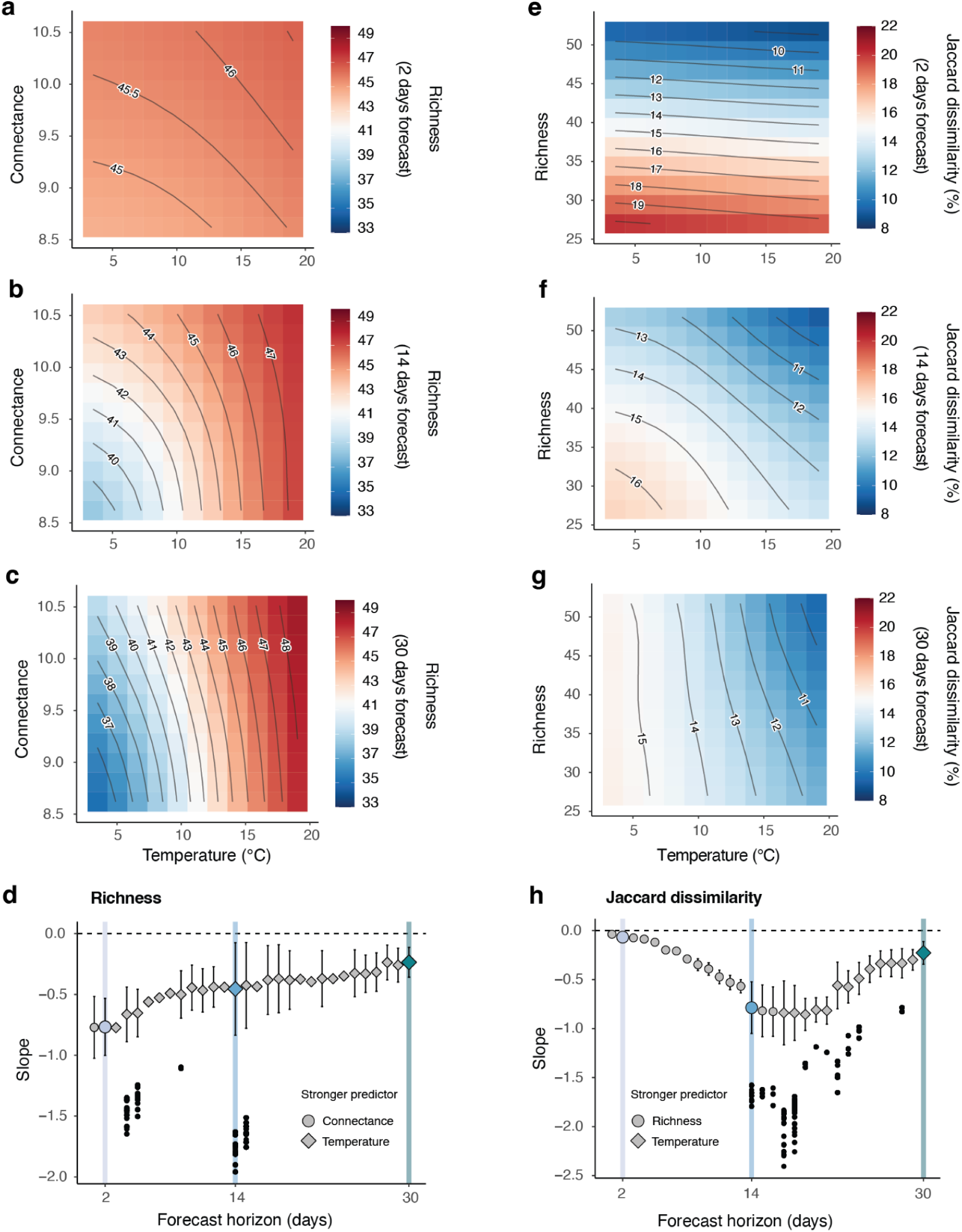
Interaction between biotic and abiotic predictors of biodiversity forecasts at different horizons. Interactive effects of water temperature and connectance on richness **[a-c]**, and of temperature and richness on Jaccard dissimilarity **[e-g]**, for 2, 14, and 30-day forecasts. The color code shows the forecasted endpoint for each value combination of the two explanatory variables on a 10 x 10 grid. Degree of nonlinearity and strength of the interaction between biotic factors and temperature on richness **[d]** and Jaccard dissimilarity **[h]** over different forecast horizons. Negative slopes indicate that the 2 predictors affect the response variable synergistically, and the larger the absolute slope, the stronger the effect (for more information, see **Fig. S5**). The error bars show the standard deviation of slopes in the 10 x 10 grid (n=100): a larger standard deviation corresponds to a stronger nonlinearity of the slope in the entire grid. The symbol represents the strongest predictors at that forecast horizon. The black dots show outliers (> 3 times the standard deviation).

Overall, the degree of nonlinearity and the strength of the interaction between temperature and biotic factors peak in short and medium-term forecasting for richness and Jaccard dissimilarity, respectively (**Fig. 4d-h**). In both cases, biotic factors (network connectance and taxonomic richness) strongly influence the interaction in short-term forecasts, while temperature takes the lead in long-term forecasting. This evidence underscores the importance of incorporating temperature values from climate projections into biodiversity forecasting models.

## Conclusions

Our results show that we can successfully forecast presence-absence-based biodiversity metrics, identify the most important biotic and abiotic predictors, and study their nonlinear interactions in a complex and natural ecosystem. There is an apparent research gap in quantitatively forecasting biodiversity metrics, especially in natural ecosystems that exhibit chaotic dynamics (i.e., plankton), where SDMs may perform poorly^59^. If we mischaracterise chaotic dynamics as noise under the assumption of equilibrium, short-term forecasts may be less accurate, and long-term forecasts may be overconfident^20^. This lack of forecast proficiency can result in unanticipated collapses of communities or events of dominance (e.g. blooms) due to the underestimated effect of human-made disturbances, which is especially problematic in short-lived organisms with rapid changes (i.e., plankton)^20^. A promising way to increase forecast proficiency in natural communities is to rely on approaches that accommodate nonlinear chaotic dynamics (e.g., methods based on state-space reconstruction). Those approaches often require data with the right information content and appropriate temporal frequency. As forecast accuracy diminishes with the forecast horizon, information on biotic interactions and continuous monitoring programs become indispensable for achieving the highest possible proficiency of forecasts. We show that biotic and abiotic predictors are equally important for forecasting biodiversity metrics in plankton communities. Therefore, monitoring emergent properties of complex ecosystems (e.g., biotic interactions) and relevant external abiotic factors (e.g., temperature) is essential for anticipating undesirable changes in biodiversity. With the decreasing costs of acquiring and processing monitoring data and the advent of automated monitoring tools, anticipatory forecasts like the one presented here could become more widespread across many complex ecosystems. In lakes, the current advancements in developing future warming scenarios could significantly enhance our ability to forecast biodiversity metrics if combined with observational time-series data that provide biotic factors. This development will be necessary to provide the tools for effective decision-making and conservation strategies to face rapid changes in ecosystem services worldwide.

## Methods

### Data collection

We obtained high-frequency data on physical and chemical water parameters, plankton abundance, nutrients, and weather from an autonomous lake monitoring platform (**Fig. S1 - Fig. S3**)^60,61^. This platform is in the middle of Lake Greifen, a small peri-alpine eutrophic lake in central Switzerland.

### Plankton abundance

An automated underwater dual-magnification microscope attached to the monitoring platform collects data on the lake’s plankton community^61^. The two magnifications allow the camera to simultaneously image three levels of the food web (primary producers, grazers, and carnivores). Plankton images were taken for 10 minutes at the beginning of every hour for four consecutive years. The images were classified into 69 taxa using a neural network, with an average accuracy of 0.9025 and F-score of 0.8838 for the higher magnification (phytoplankton) and an average accuracy of 0.9376 and F-score of 0.8956 for the lower magnification (mostly zooplankton)^62^, https://github.com/kspruthviraj/Plankiformer). We subset the images of the lower magnification to every 6 seconds to exclude multiple imaging of the same object since some individuals were observed to remain in the camera’s focal plane for some seconds. We aggregated (sum) images per taxon over one day (00:00-23:59) to account for daily fluctuations in plankton abundance and reduce observational noise. We converted the number of images per second taken (ROI=raw object of interest per second) into indv./mL based on laboratory calibrations to compare the two magnifications ^61^. Over the four years, we collected 1382 observations and excluded days that did not have at least 20 hours sampled because plankton displays strong fluctuations during the day (e.g., daily migration of zooplankton^63^, and our samples would be biased. After excluding the incomplete samples (n=50), we had 1332 remaining sampling dates. Because the analysis methods are sensitive to rare taxa, we excluded taxa that did not occur over 50% of the time (positive abundance on a given day) and kept 55 taxa for further analysis ^27^ (**Tab. S1, Tab. S2**).

### Biodiversity metrics

Richness or alpha-biodiversity counts the number of different species in a community. A community is limited by an upper number of species (taxa), depending on resources and the environmental context. The more taxa there are, the slower the expected turnover in biodiversity. Shannon diversity is calculated based on the number of species and their relative abundances. The index is greater when more unique species are in the community or when species evenness is high. Shannon diversity considers taxa abundances, while richness is based on presence-absence only.

We use Jaccard and Bray-Curtis dissimilarity (**Fig. S2**) as indicators of change in plankton community composition, an essential aspect of biodiversity. Bray-Curtis dissimilarity considers changes in taxa abundances, while Jaccard dissimilarity is based on presence-absence only.

### Temperature

Temperature is a fundamental driver of taxa’s physiological processes (e.g., metabolism, respiration, photosynthesis, or activity patterns), influencing how biomass is distributed between taxa and the ecosystem^64–66^. Moreover, the water temperature can affect a lake’s water column stability and, consequently, water oxygen and phosphorus levels^67,68^. In the past decade, the water temperature in lakes has been rising alarmingly, and it is expected to alter aquatic ecosystems fundamentally^31,69,70^. Water temperature was measured with a CTD probe from the surface to the lake’s bottom (1-17.5 m) at the monitoring platform and then averaged over the water column.

### Light

Light penetrating the water column allows primary producers such as phytoplankton to do photosynthesis, allowing them to grow and make biomass available for higher trophic levels. Moreover, it represents how much energy the system receives from the sun. We used light penetration to measure the light available in the water column. We calculated light penetration by defining the depth at which PAR (irradiance) measured by a CTD probe reaches 5, the lowest detectable value. PAR is a measurement of light emission within the range organisms can use for photosynthesis (400-700 nm). We choose light penetration instead of PAR (irradiance) as a measure of light to avoid a bias introduced by the monitoring platform shading the CTD probe in the upper part of the water column.

### Wind and mixed layer depth

Disturbances of the water column can alter the abiotic environment and drastically reshuffle the plankton community, leading to a turnover in taxa. We used the maximum wind speed and the depth of the mixed layer as indicators of the water column’s physical disturbance. Moreover, strong winds can push the thermocline downwards, enabling the exchange of heat, oxygen, and nutrients between bottom and surface waters. The wind speed was estimated using a local weather station attached to the monitoring station’s roof. We used the maximum wind speed reached for the day. Seasonal mixing, expressed as the depth of the mixed layer, makes nutrients from the lake bottom available to organisms living in the photosynthetic zone (upper 8 m), triggering the growth of primary producers and making biomass available for upper levels of the food web. The mixed layer is expected to be deepest in winter and spring when the temperature in the lake is almost constant across the depth and thus, the water column is thoroughly mixed. In summer, the lake is fully stratified due to strong temperature gradients in the water column, separating the nutrient-rich oxygen-poor cold bottom water from the nutrient-poor oxygen-rich warm surface water. We calculated the mixed layer’s depth based on a CTD-probes temperature profile and the R package rLakeAnalyzer ^71^.

### Nutrients

Phosphate (PO_4_), nitrate (NO_3_), and ammonium (NH_4_) are essential nutrients for the growth of plankton. Primary producers, e.g., phytoplankton, can uptake them directly from the environment, while higher taxonomic levels, such as grazers, obtain nutrients through the food chain. Organisms use phosphorus in fundamental processes such as the storage and transfer of genetic information, cell metabolism, and the energy system of the cells. Phosphorus is taken up by organisms mostly as inorganic phosphate (PO_4_) and is the main limiting nutrient for lake productivity. Nitrogen may become the main limiting factor for algal growth in lakes with very high phosphate concentrations. Ammonium (NH_4_) is the energetically preferred nitrogen source since a cell must reduce nitrate (NO_3_) before assimilating it. We measured phosphate, nitrate, and ammonium weekly next to the monitoring platform. We interpolated the weekly nutrient time series to obtain daily data using a trained random forest model based on CTD-probe and weather station data. For the interpolation, we used for all nutrients: water temperature (mean, photic), conductivity (mean, photic), oxygen concentration (mean, photic), oxygen parts per million (mean, photic), pH (mean, photic), as explanatory variables. In addition, we used air pressure (mean, absolute) as an explanatory variable for phosphate, depth at 5 PAR (see “Light” section) for nitrate, and air temperature (mean) for ammonium. The model performance (R^2^) was 0.90 for phosphate, 0.77 for nitrate, and 0.70 for ammonium. We used the randomForest package (version 4.6-14) in R to train and run the random forest models. We predicted nutrients at the frequency of the CTD data (every 3 hours in summer, 6 hours in winter) and averaged them over a day.

### Trace of the interaction matrix

We want to test the forecast ability and effect on biodiversity change of an important biotic variable, the trace of the Jacobian interaction matrix (sum of the first diagonal). The trace provides information on a community’s sensitivity to external perturbations (i.e., in its environment) by measuring the divergence of future trajectories of a system (greater divergence = more unstable)^30^. It combines and summarises species interdependencies (biotic interactions) and single-species sensitivities to environmental change (intrinsic growth). Moreover, the trace has been linked to the structural stability of a community and forecast ability (high trace = structurally unstable and harder to forecast). We can estimate the trace from chaotic nonlinear dynamics, and it does not require the system to have an equilibrium. Plankton exhibits chaotic nonlinear dynamics with no equilibrium, and the underlying equations are hard to unravel with noisy observational data. Moreover, the trace is more robust to observational noise than estimating community stability based on dominant eigenvalues or Lyapunov exponents^30,72,73^. In our work, we link community sensitivity to a change in biodiversity, i.e., community composition (Bray-Curtis and Jaccard dissimilarity and the gain and loss of taxa).

### Connectance

Connectance quantifies the percentage of realized links (interactions) compared to all possible links at a given time point in a community. We considered a link between two species if the time-varying Jacobian interaction coefficient differed from zero. Connectance can increase robustness in food webs and has been positively linked to linear stability in theoretical ecology^74^.

### Interaction coefficients (Jacobian matrix)

We used the multiview distance S-map method (MVD S-map) from the EDM framework to estimate the time-varying interaction matrix in the plankton community^27^. S-map models are local multilinear regressions weighted based on state space proximity^22^. By averaging the distance matrix from multiple smaller embeddings than the community size, MVD S-maps are more robust towards the problem of overfitting than the original S-map. This addition allows the reconstruction of interaction networks of n-dimensional communities. Moreover, by including regularization schemes, it can deal better with observational noise and collinearity among model parameters ^26^. To do so, we had to i) calculate the distance between sampling points ii) parametrize the local regularized regression, and iii) extract the local model coefficients. The local model coefficients are an approximation of the Jacobian interaction matrix.

We scaled the taxa time series for zero means and standard deviation before reconstructing time-varying networks with the MVD Smap analysis using the scale function in R.

#### Distance of sampling points in state space

We weighted the data points according to their position in state space before fitting the regression to account for nonlinear dynamics. We calculated the Euclidean distance among points in low dimensional state space reconstructions (= multivariate embeddings) and took a weighted average across all multivariate embeddings. Calculating the distance in multiple low-dimensional embeddings allows us to overcome the curse of dimensionality^27^. We had to select the multivariate embeddings with the highest predictive power. This step was necessary because of computational limitations.

##### Multivariate simplex projection

We used the target taxon and 9 other taxa (E = 10) to create different possible multivariate embeddings. Because of computational limitations, we set the limit of the maximum number of different multivariate embeddings to 1’000. For taxa, where we could build more than 1’000 embeddings, we subsampled 1’000 randomly. We estimated the one-day-ahead (tp = 1) forecast skill of those multivariate embeddings using simplex projection on our target taxa. We chose the top 100 embeddings based on minimizing forecast error (rmse). Because of high autocorrelation, we excluded the 14 closest points in attractor space from predictions (exclusion radius = 14)

##### Multiview distance

We then calculated the “multiview distance” (average distance between points) across the top 100 embeddings. To do so, we calculated the Euclidean distance among points in the state space for each multivariate embedding. Then we calculated a weighted average across all embeddings of the estimated distance between sampling points in attractor space, resulting in the “multiview distance”. The chosen weights were based on the forecast error rmse previously estimated, where multiview embeddings with smaller rmse had a higher weight. Those distances later served as weights in the MVD S-map.

#### Local linear regressions (S-map)

We used the coefficients of a local elastic-net regression weighted in state space to approximate the time-varying interaction strength between taxa and reconstructed the local interaction matrices. Elastic-net regression uses a mixture of lasso and ridge penalty. Regularization techniques allow the model to generalize and improve its performance on unseen data. They make the model simpler by reducing the magnitude of the coefficients. They can be used to prevent under- and overfitting. Moreover, they help cope with observational noise in the data.

##### Parametrization

We had to parametrize the local elastic-net regressions before extracting the interaction coefficient. In detail, we had to parametrize theta (controlling for the model’s degree of nonlinearity, high theta = nonlinear; theta zero = linear), alpha (balancing between ridge and lasso regression penalty, 0.5 we use an equal mix of both, whereas with 1 we exclusively use lasso penalty and 0 ridge penalty) and lambda (controls the strength of the penalty, large lambda = high penalty, pushes most coefficients towards or close to 0) for the MVD S-map. We used all possible combinations (n=125) of theta (0, 0.1, 1, 3, 8), alpha (0.1, 0.3, 0.5, 0.7, 0.9) and lambda (1.000, 0.178, 0.032, 0.006, 0.001) to evaluate the predictive performance of MVD S-map. When longer time series are available, explicit training and testing data sets are more robust than leave-one-out cross-validation ^73^. Moreover, time series have temporal dependencies that must be preserved during cross-validation (i.e., we cannot use future points to predict past values). Therefore, we parametrized the regularized s-maps using “day-forward-chaining” to overcome this issue. The initial library contains the first 10% (n = 139) data points, which are used to predict the next point (here, t =140). This point is then added to the library to predict the next point. With this approach, we predicted the remaining 90% of the data, always increasing our library size with the predicted value. We evaluated the predictive performance of the MVD S-maps using the rooted mean squared error (RMSE).

##### Model averaging

Many models (parameter combinations) have similar forecast errors but differ substantially in estimating the interaction coefficients. Thus, we select the top 5% (n=6) models based on the lowest mean squared error (rmse). We take a weighted average of the coefficients estimated by the top models to estimate interaction with higher accuracy (Cenci and Saavedra 2018). Moreover, the standard deviation and coefficient of variation of the coefficients allow us to assess their variability and uncertainty.

#### Differences between our approach and the original MVD S-map

We simplified the original MVD S-map approach proposed by^27^ to make it less computationally intensive and conceptually easier while maintaining the same efficacy (**Fig. S4**). In detail, we:

1. Take E=10 for all taxa - we do not do a univariate simplex projection to find the best E, since many possible E result in very similar predictive power in our case.
2. include all taxa in the multivariate embeddings - we do not do ccm to find the most important interactions, since this does not improve the forecast skill of the model and is computationally intensive; this also means we did not consider more than 1 lag; moreover, this pre-selection of interactions only applies to the MVD distance calculations, for the final S-map models all taxa are used.
3. Use 1000 multivariate embeddings instead of 10,000—this reduces computation time, and there is no significant improvement by having more than 10,000 multivariate embeddings.
4. We select the top 100 multivariate embeddings based on rmse for consistency (instead of rho).
5. We use day-forward-chaining cross-validation to parametrize the MVD S-maps (instead of leave-one-out cross-validation since this is more appropriate for long-time series data). An illustrative explanation of this approach can be found in the following blog post^1^.
6. Only use 125 parameter combinations of alpha, theta and lambda to parametrize dem MVD S-maps - instead of 2025, there is no major improvement in inferring interaction coefficients with more than 125 parameter combinations and it drastically reduces the needed computational resources.
7. Finally, we averaged the model coefficients for the top 5% (n = 6) MVD S-map models - since different parameter combinations yielded very similar forecast performance and coefficients can vary significantly depending on the model parametrization.

To ensure that our simplified approach does not reduce the MVD S-map efficacy in estimating network properties, we used the same theoretical population model (Multispecies Ricker Model) that has been used to validate the original approach^27^. We calculated network properties based on the coefficients provided in the **Supplementary** of Chang et al. 2021 and used the Multispecies Ricker model time series provided in the **Supplementary** to run our simplified approach. The scaled trace of the theoretical Jacobian correlated well with the approximated trace by MVD S-map for both the original and simplified approach (**Fig. S4a**). Connectance was constant in the theoretical model, and approximated connectance by MVD S-maps only differed slightly from a constant value for both the original and simplified approach (**Fig. S4b**).

### Forecasting biodiversity metrics

Biotic and abiotic mechanisms such as taxa interactions and temperature variation can affect plankton biodiversity, i.e., lead to a change in richness, diversity, or turnover in community composition. We used regularized S-maps (elastic-net) to forecast biodiversity based on abiotic and biotic variables. In detail, we forecasted richness, diversity, Bray-Curtis and Jaccard dissimilarity 1 to 30 days ahead (this forecast horizon encompasses 3-30 generations of our focal organisms). We used a separate forecast model for each horizon (**Fig. S5a**). For all biodiversity target variables, we use connectance, the trace of the interaction matrix, water temperature, phosphate, ammonium, nitrate, maximum wind speed, mixed layer depth and light penetration as explanatory variables. For the Jaccard dissimilarity, we included richness; for Bray-Curtis dissimilarity, we included diversity as additional explanatory variables. Moreover, each model included an autoregressive term (i.e., the biodiversity target variable) to control for potential autocorrelation. The resulting model formulas were:

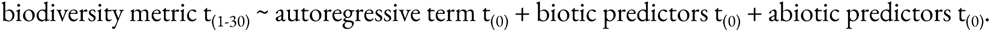

We parameterized the regularized S-maps (elastic-net) for each forecast horizon analogous to the MVD S-map to infer the taxa interaction coefficients (based on day-forward-chaining cross-validation and 125 parameter combinations for alpha, theta and lambda) (**Fig. S5b-c**). Estimating the MVD was unnecessary (we have only ten or eleven explanatory variables). Thus, we estimated the distance among points in SSR based on the ten or eleven explanatory variables. Again, we kept the top 5% parameter combinations (n = 6) for each model - our forecast uncertainty estimates come from the proficiency of these 6 models. This resulted in 6 models per forecast horizon and biodiversity metric, resulting in a total of 720 separate models (**Fig. S5d**).

### Forecast proficiency

We calculated forecast proficiency based on skill (Pearson’s correlation between observations and predictions) and error (rooted mean squared difference between observations and predictions) from the cross-validation and the final model fit for each horizon.

### Model coefficients

We estimated the coefficients from the best-fitted models (using all available observations after cross-validation) and averaged them (weighted based on the lowest rmse) to examine the direction and magnitude of each predictor’s effect on biodiversity metrics. The larger the absolute values of the S-map model coefficients, the stronger the effect of the explanatory variable (biotic and abiotic predictors) on the target response variable (biodiversity metric). The sign of the coefficient indicates the direction of effect: positive for positive correlation, negative for negative correlation. Coefficients close to zero indicate that mechanisms are not relevant to the model. Furthermore, the coefficients of the S-map can be used to rank the mechanisms driving biodiversity turnover by their importance.

### Variable importance

We estimated the importance of each variable for forecasting biodiversity by fitting a model without this variable (including the day-forward changing cross-validation). We then compared the forecast skill (rho) and forecast error (rmse) from the model with and without that variable.

### Interactions between biodiversity drivers

We want to test how the interaction between biotic and strong abiotic factors can affect plankton biodiversity change. Therefore, we predicted the response variable (richness and Jaccard dissimilarity) over a varying pair (=gradient) of biotic and abiotic explanatory variables. We choose connectance and water temperature as explanatory variables for predicting richness, and richness and water temperature as explanatory variables for predicting Jaccard dissimilarity. We varied the explanatory variables from the minimum to the maximum observed value with ten regular intervals (resolution), resulting in 121 explanatory variable pairs. We predicted the response (change in biodiversity) up to 30 days ahead for those variable pairs using the best previously parametrized regularized S-map model. We did this for each time point by setting the remaining variables at their current value. Setting the remaining variables at their current value creates a more realistic scenario than keeping them at their median, which can result in combinations rarely or never observed in the lake. To reduce the computation power required, one could predict fewer time points by randomly subsampling the time series.

We estimated the interaction between two explanatory variables by estimating the slope of the contour lines. To do so, we extracted the contour lines with a resolution of 10 x 10 points (going from the minimum to the maximum observed value, evenly spaced). We approximated the contour line for each grid cell using a linear model and extracted the slope. To avoid slopes going to infinity, we estimated the slope in both directions (y ∼ x, x ∼ y) and kept the smaller slope. We then averaged the slopes across the whole grid and calculated the standard deviation. A mean slope close to zero or one indicates that one variable dominates the relationship (= has a stronger effect on the response variable) and there is no interaction between the two explanatory variables (**Fig. S6 a-b**). A negative slope indicates a synergistic interaction (= additive effect) between the two explanatory variables (**Fig. S6 c**) and a positive slope indicates an antagonistic interaction (= explanatory variables cancel each other out) (**Fig. S6 e**). A large standard deviation indicates a nonlinear relationship between the two explanatory variables (many different slopes), indicating that the interaction is state-dependent (**Fig. S6 d,f**).

---

**Fig. E1:**
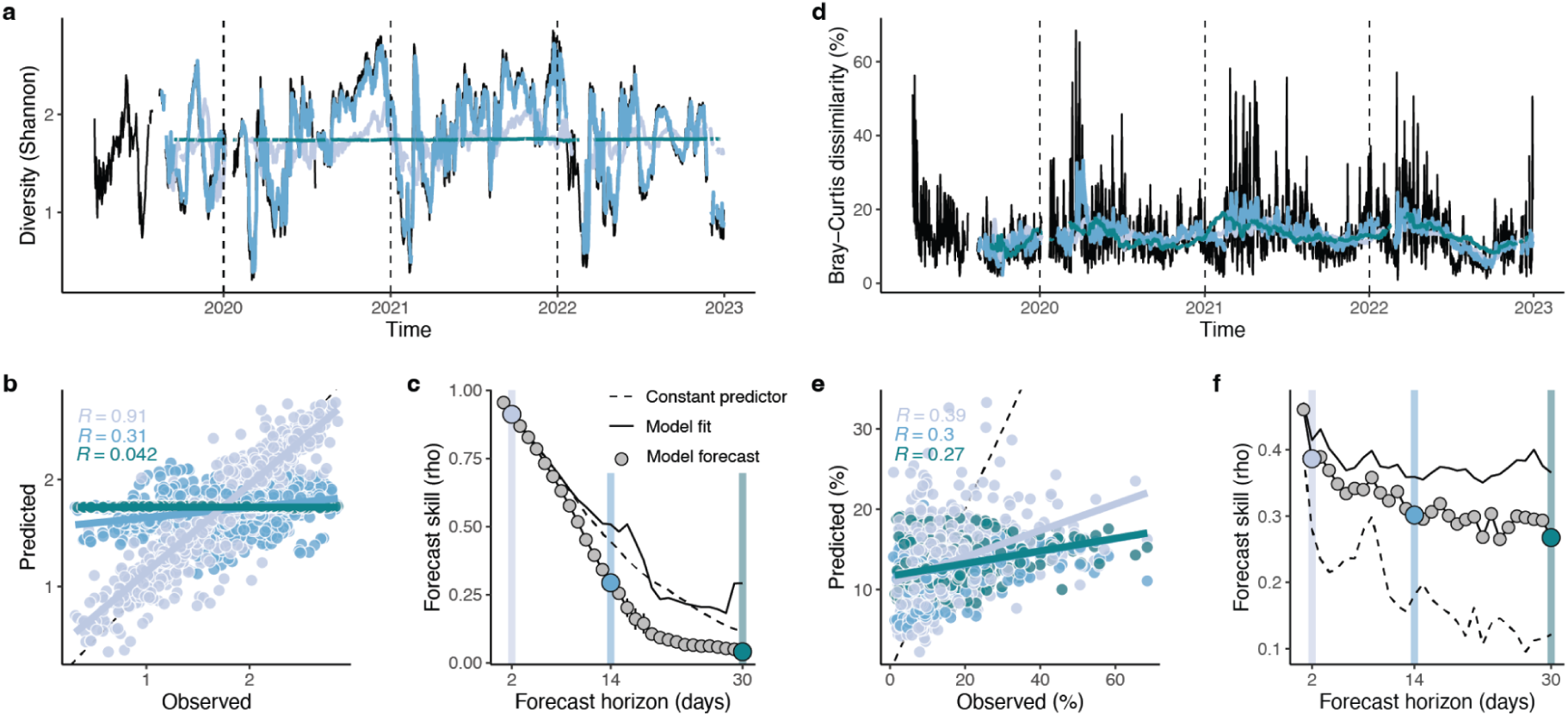
Forecast proficiency when predicting biodiversity change (Shannon and Bray-Curtis dissimilarity) based on biotic and abiotic factors. **[a** and **d]** Time series of observed and predicted values for the forecast horizons 2, 14 and 30 days. **[b** and **e]** Correlation between predicted and observed values for Shannon and Bray-Curtis dissimilarity, respectively: the dotted line indicates the 1 to 1 line. **[c** and **f]** Forecast skill (estimated as the Pearson correlation coefficient between observations and predictions) of the six best models (average ± standard deviation) for Shannon and Bray-Curtis dissimilarity, respectively.

**Fig. E2:**
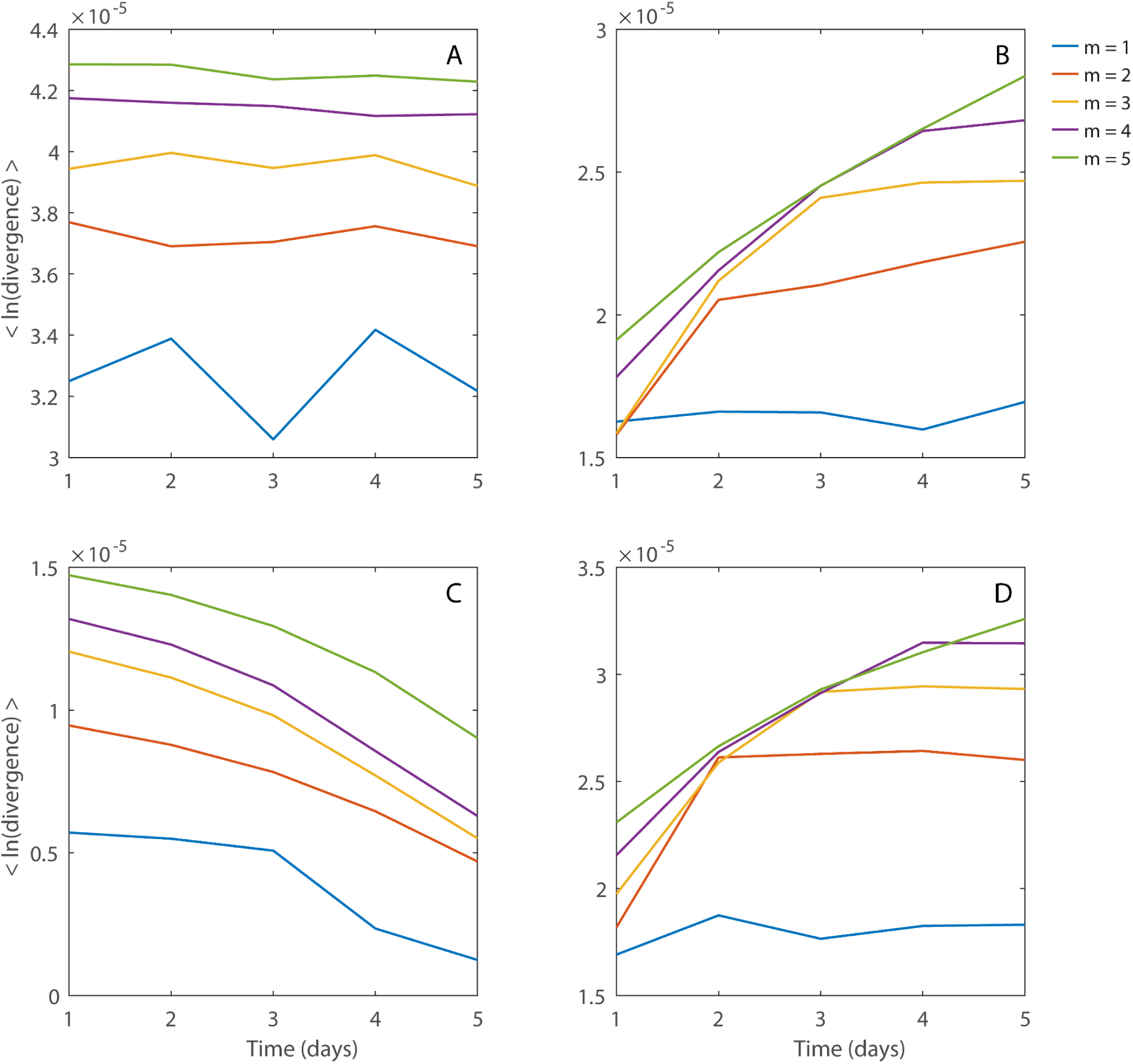
Diagnosing chaos in biodiversity metrics. Trajectory divergence as a function of time for different embedding dimensions (m) following^1^ for the time series of A) Species richness, B) Jaccard dissimilarity, C) Shannon diversity, and D) Bray-Curtis dissimilarity. Since the algorithm requires an equally-sampled time series to calculate divergence, we used the longest stretch of the time series that contains no gaps, which is 577 days. We do not provide an exact estimation of the Lyapunov exponent, since diagnosing chaos doesn’t require a precise estimation of the exponent—for which we would have to meet the Eckmann-Ruelle criterion^2^. Following ^1^, the clear presence of a positive slope, and the convergence as the embedding dimension increases, indicate chaos in the time series. Species richness (A) and Shannon diversity (B) do not follow chaotic dynamics. However, Jaccard (C) and Bray-Curtis dissimilarity (D) follow chaotic dynamics.

**Fig. E3:**
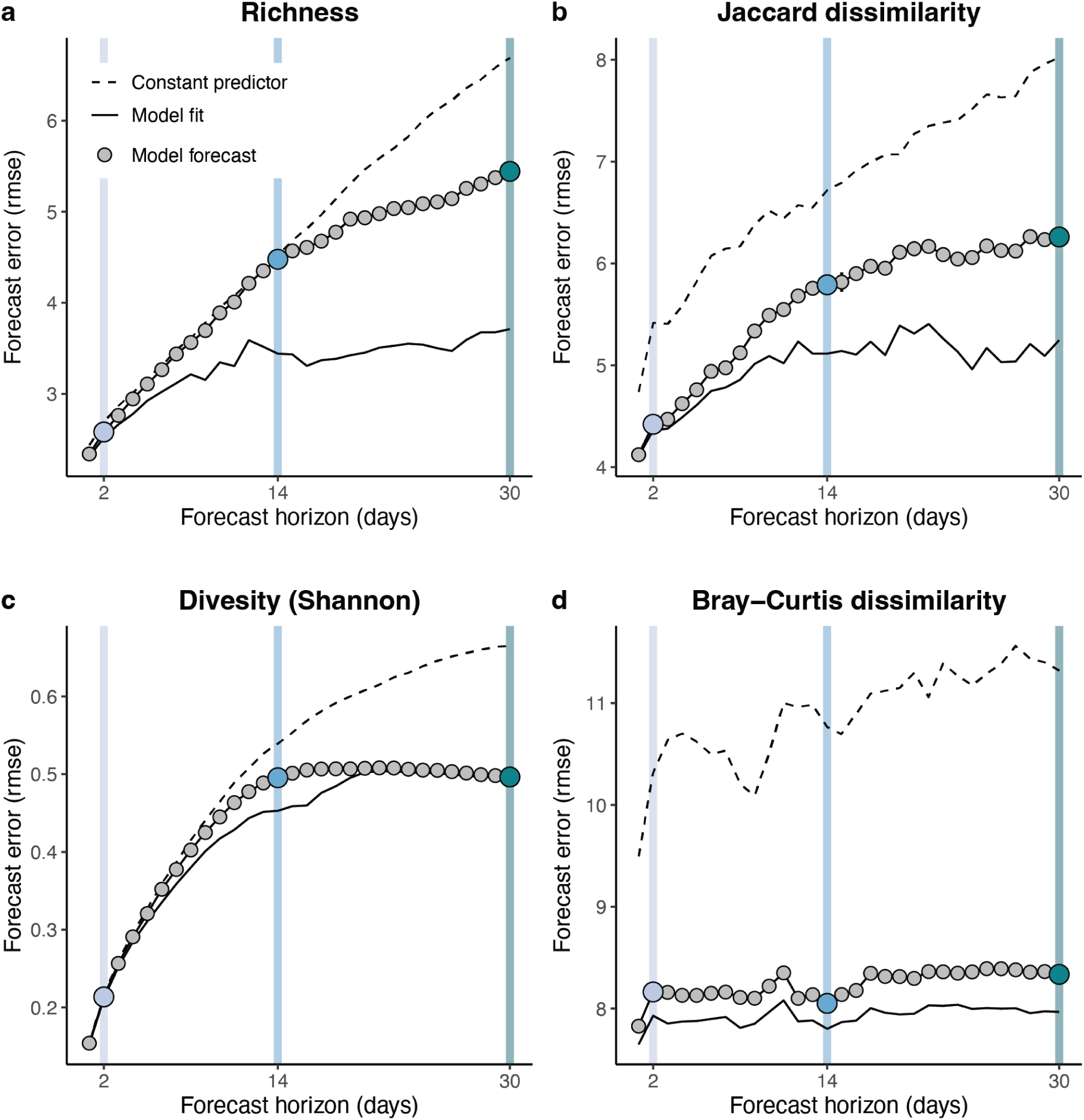
Biodiversity metric forecast error. Difference between observations and predictions, estimated as rooted mean squared error) of the six best models (average ± standard deviation) for community richness, Shannon diversity and turnover (Jaccard and Bray-Curtis dissimilarity).

**Fig. E4:**
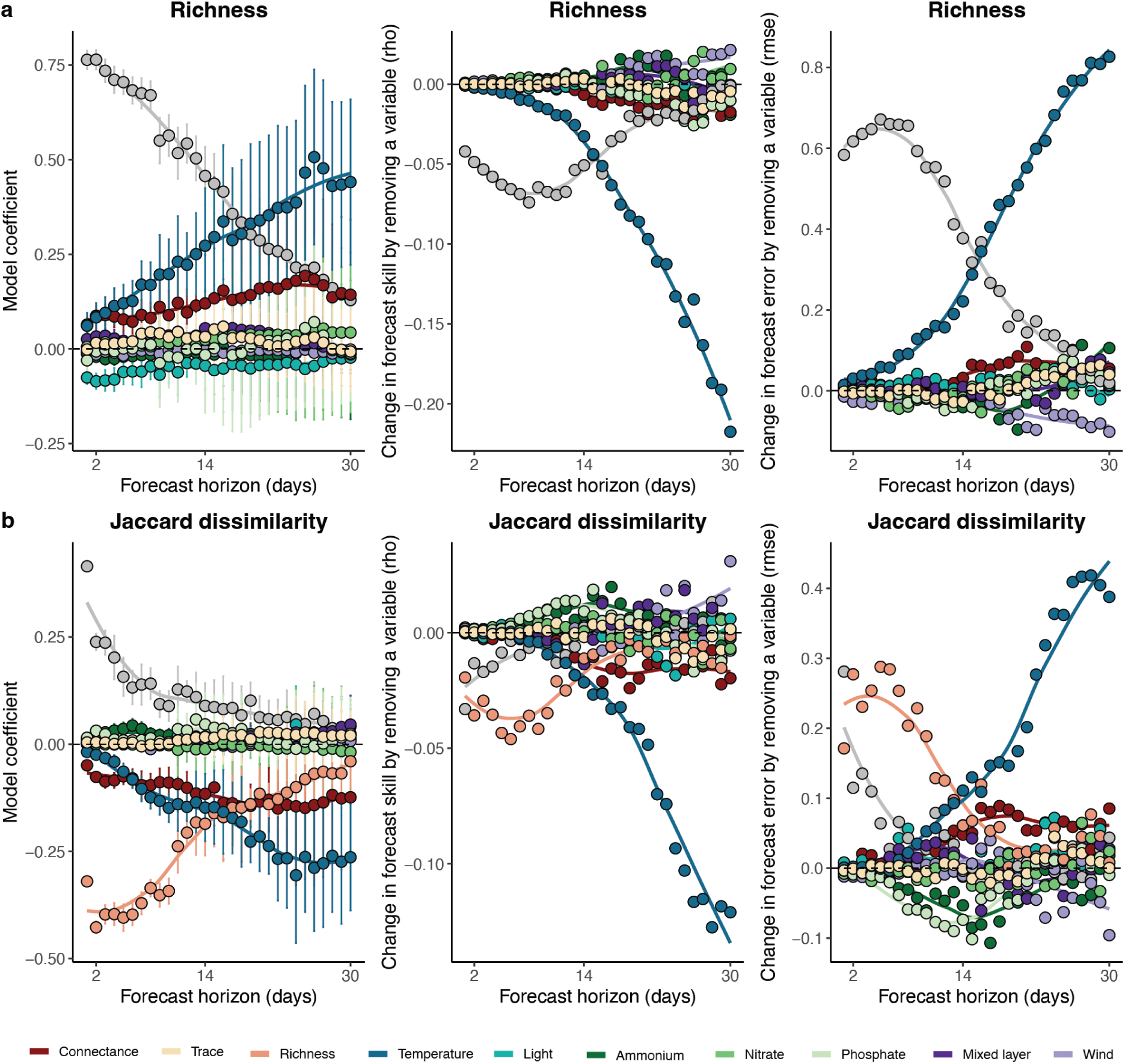
Variable importance for forecasting presence-absence-based biodiversity metrics. Variable importance can be evaluated by i) studying the model’s coefficient, where a large absolute coefficient indicates higher importance and ii) studying the change in forecast proficiency (skill and error) when omitting a variable from the forecast model. In [a] results for richness and in [b] Jaccard dissimilarity are shown.

**Fig. E5:**
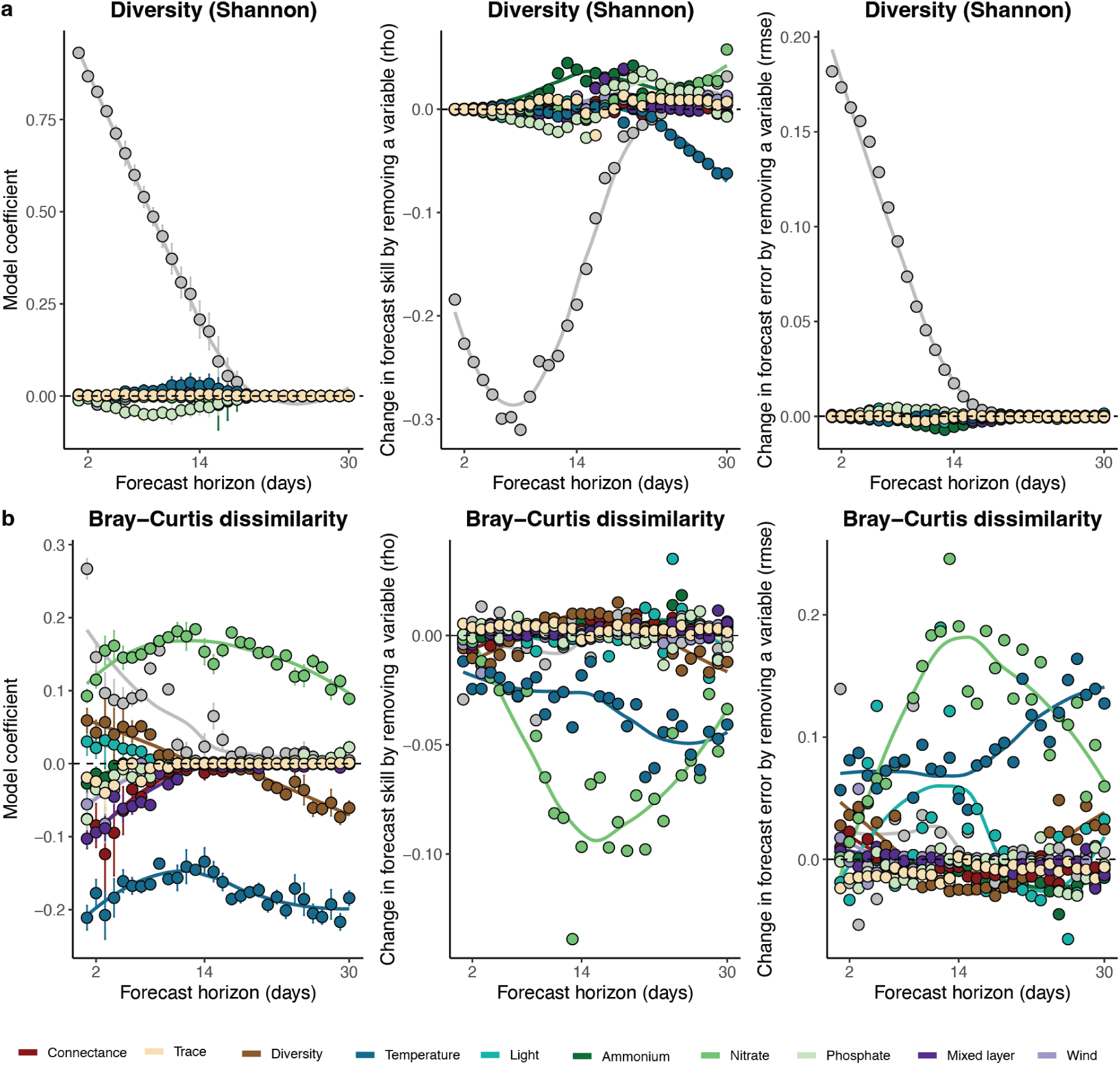
Variable importance for forecasting abundance-based biodiversity metrics. Variable importance can be evaluated by i) studying the model’s coefficient, where a large absolute coefficient indicates higher importance and ii) studying the change in forecast proficiency (skill and error) when omitting a variable from the forecast model. In [a], results for diversity and in [b] Bray-Curtis dissimilarity is shown.

## Footnotes

1 https://medium.com/@soumyachess1496/cross-validation-in-time-series-566ae4981ce4

## Notes

### Competing Interest Statement

The authors have declared no competing interest.

