## Supplementary Information for "Biodiversity forecasting in natural plankton communities reveals temperature and biotic interactions as key predictors"

*These authors contributed equally

Supplementary Figures ...................... Page 2-14

Fig. S1: **Taxa time series.**

Fig. S2: **Time series of all abiotic explanatory variables used in the study.**

Fig. S3: **Time series of all biotic responses and explanatory variables used in the study.**

Fig. S4. **Sensitivity analysis: Theoretical network properties vs. estimated by MVD S-map**

Fig. S5. **Illustration of forecasting approach.**

Fig. S6. **Interpreting the heatmap plots in Fig. 4 to understand the interaction between explanatory variables (six common cases).**

Fig. S7. **Model performance for estimating taxa interactions.**

Supplementary Tables ...................... Page 15-18

Tab. S1. **Summary statistics of taxonomic groups used for the study.**

Tab. S2. **Scientific classification of taxonomic groups used for the study.**

### Supplementary figures

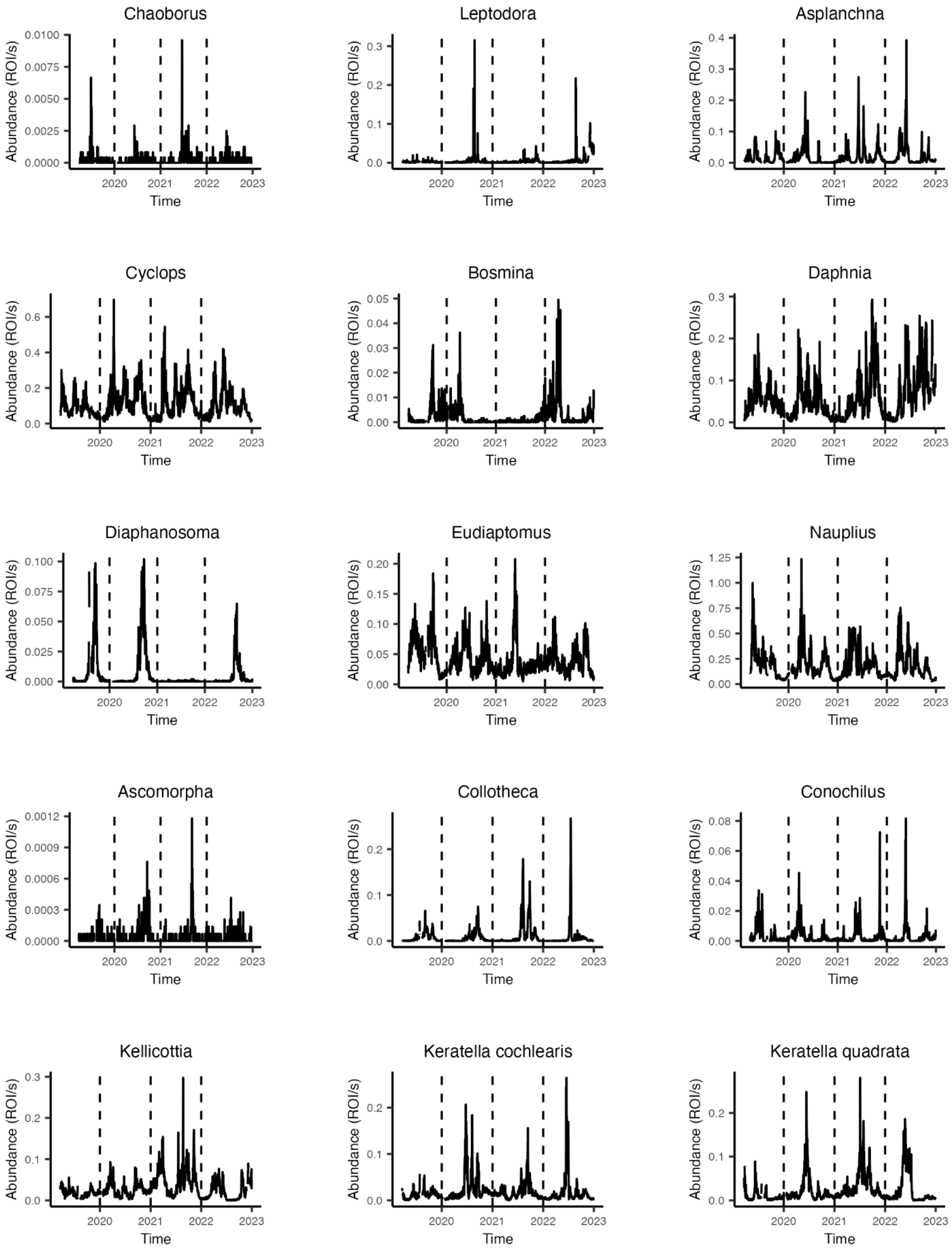

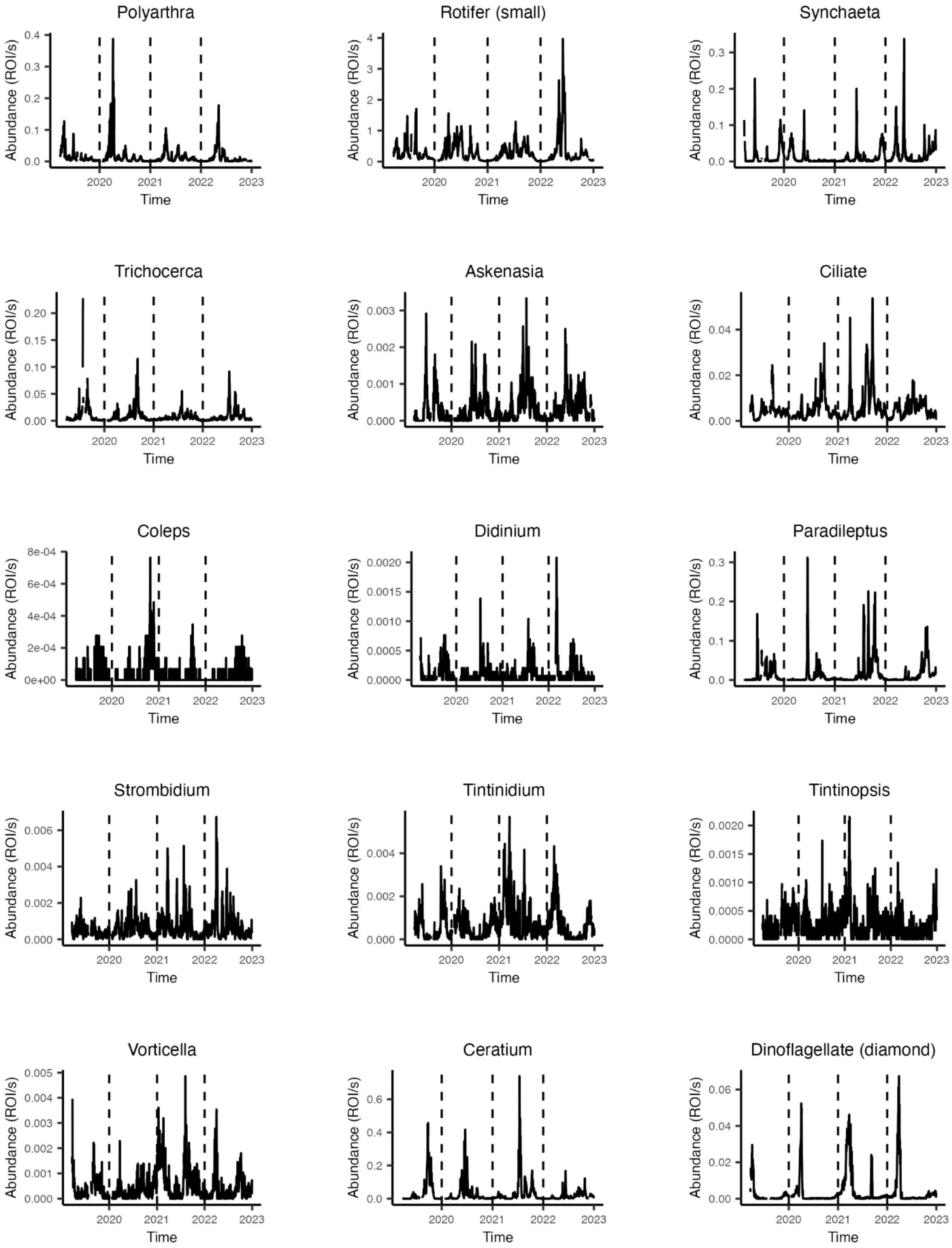
Polyarthra
Rotifer (small)
Synchaeta
Trichocerca
Askenasia
Ciliate
Coleps
Didinium
Paradileptus
Strombidium
Tintinidium
Tintinopsis
Vorticella
Ceratium
Dinoflagellate (diamond)
Abundance (ROI/s)
Time
2020
2021
2022
2023
0.0
0.1
0.2
0.3
0.4
0
1
2
3
4
0.00
0.05
0.10
0.15
0.20
0.000
0.001
0.002
0.003
0.02
0.04
0e+00
2e-04
4e-04
6e-04
8e-04
0.0000
0.0005
0.0010
0.0015
0.0020
0.004
0.006
0.005
0.6
0.06

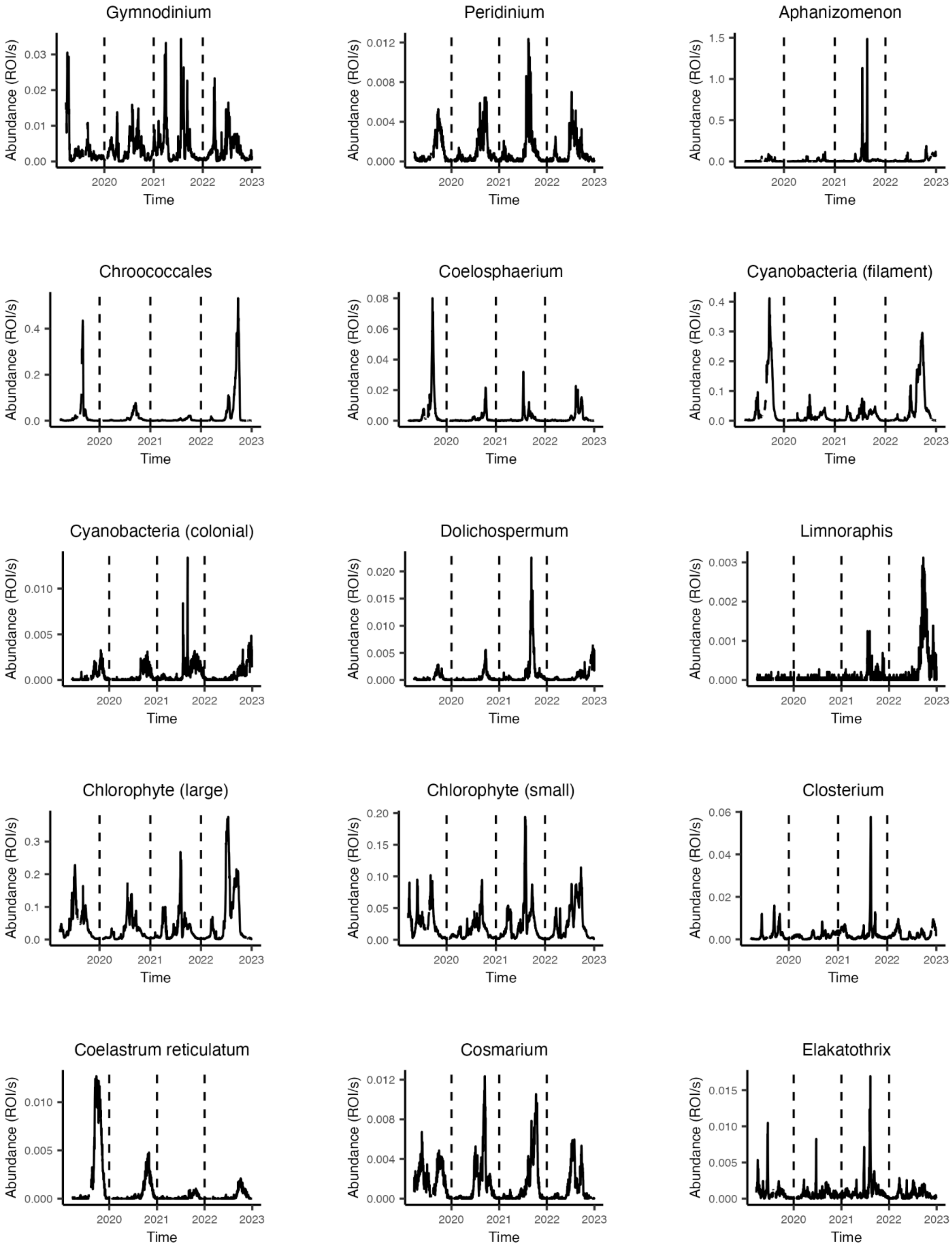
Gymnodinium
Peridinium
Aphanizomenon
Chroococcales
Coelosphaerium
Cyanobacteria (filament)
Cyanobacteria (colonial)
Dolichospermum
Limnoraphis
Chlorophyte (large)
Chlorophyte (small)
Closterium
Coelastrum reticulatum
Cosmarium
Elakatothrix
Abundance (ROI/s)
Time
2020
2021
2022
2023

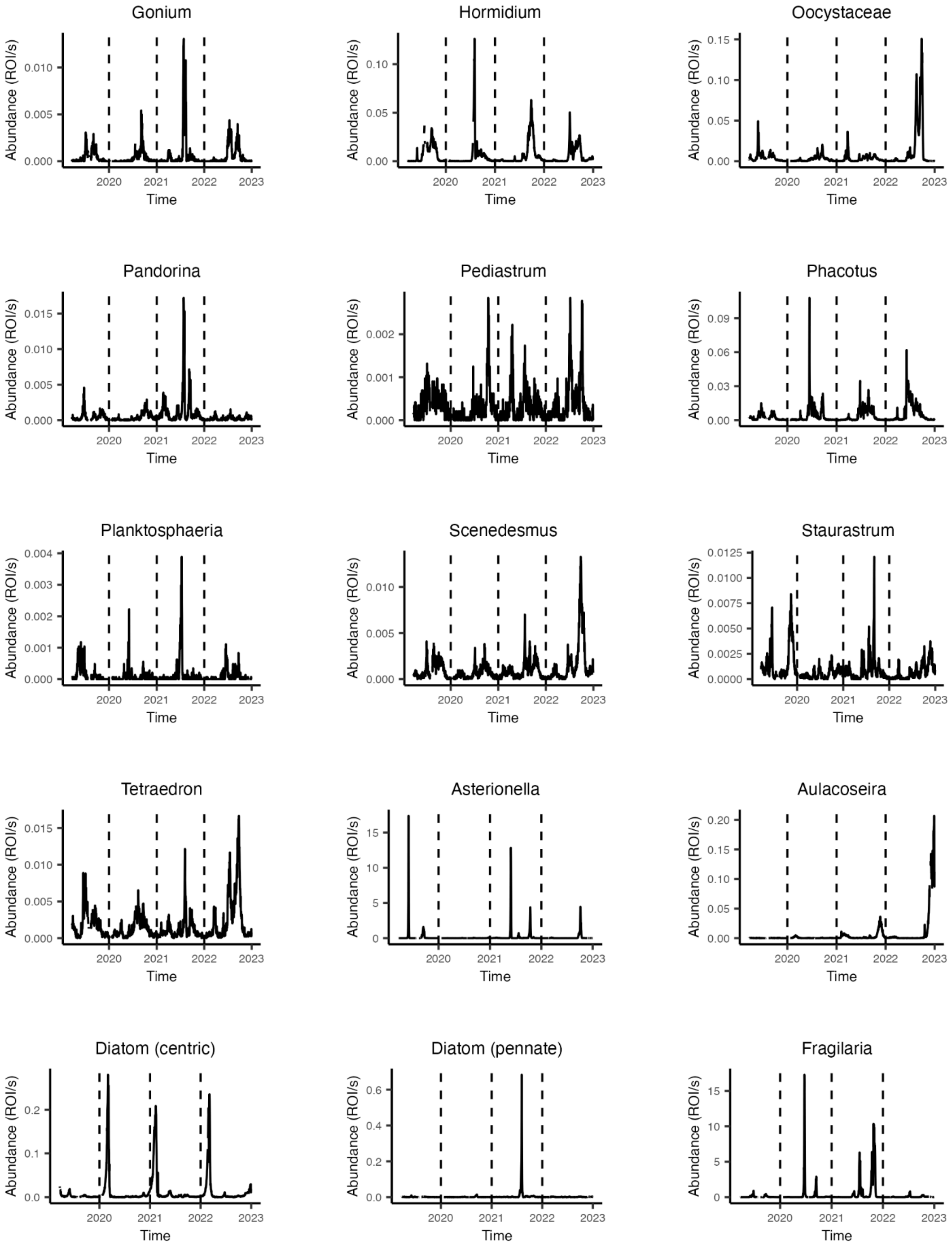
Gonium
Hormidium
Oocystaceae
Pandorina
Pediastrum
Phacotus
Planktosphaeria
Scenedesmus
Staurastrum
Tetraedron
Asterionella
Aulacoseira
Diatom (centric)
Diatom (pennate)
Fragilaria
Abundance (ROI/s)
Time
2020
2021
2022
2023

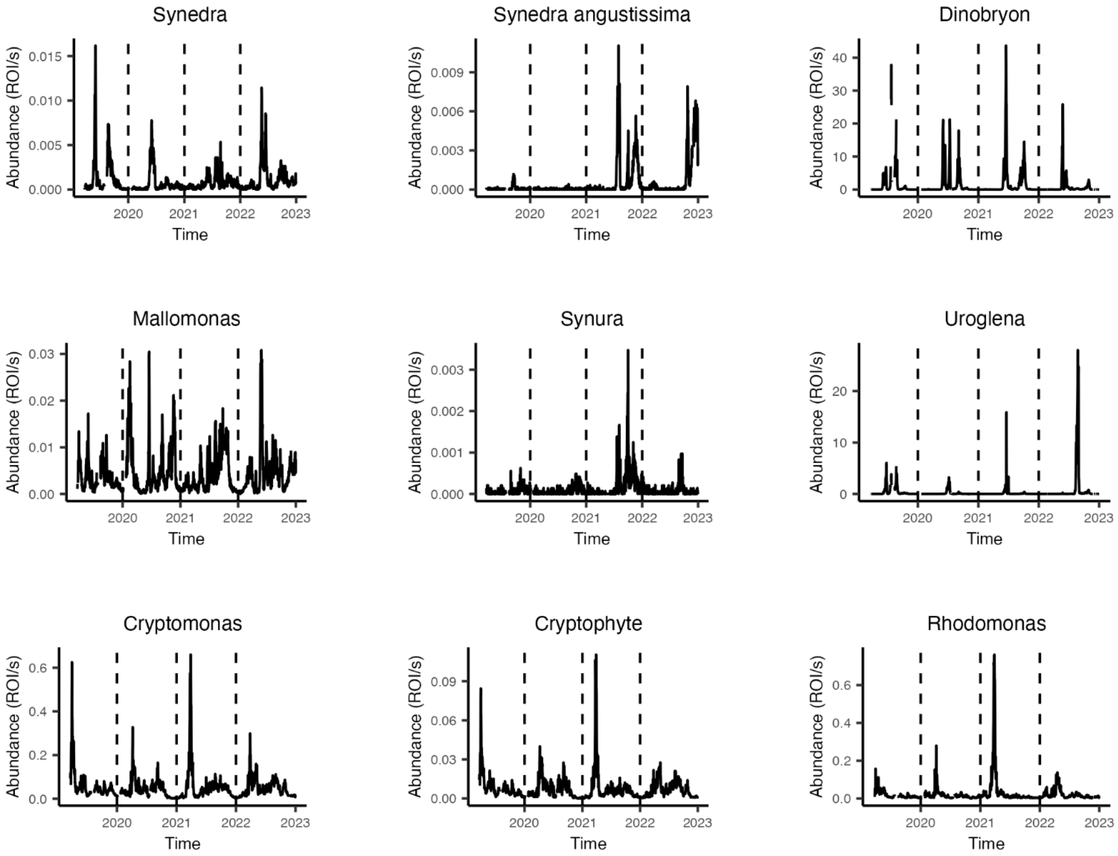


Fig. S1: **Taxa time series.** Abundances are in ROI (raw object of interest) per second.

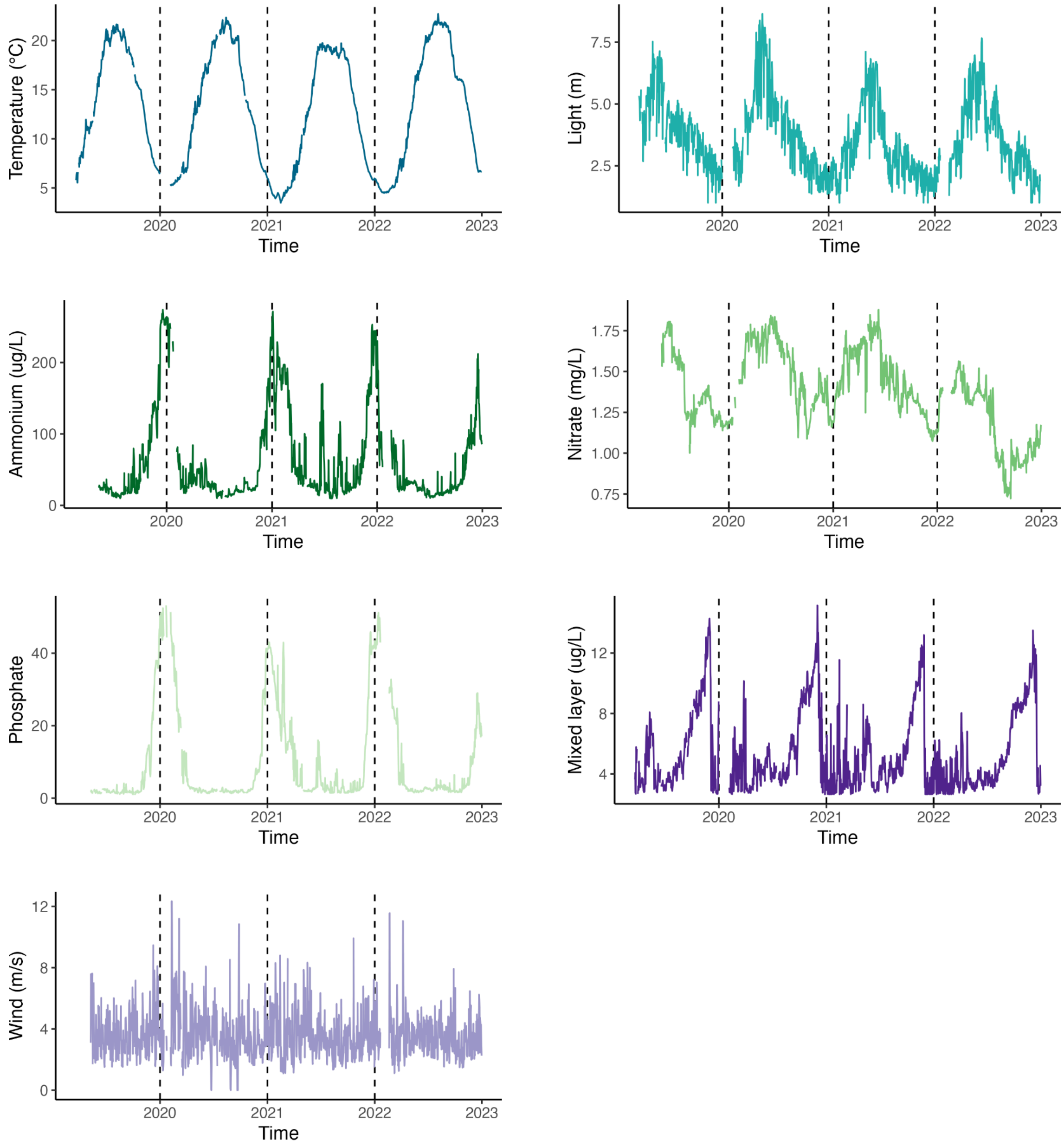


Fig. S2: **Time series of all abiotic explanatory variables used in the study.**

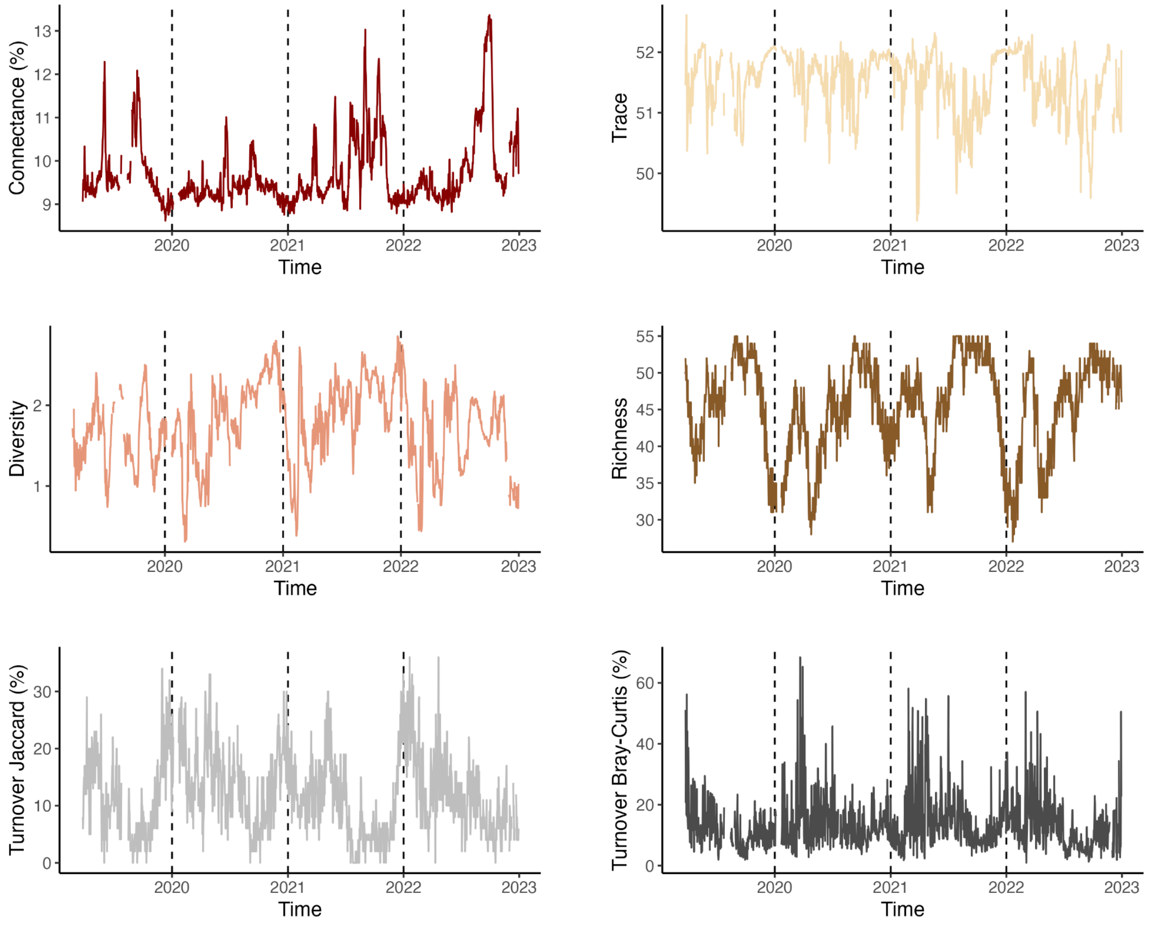


Fig. S3: **Time series of all biotic responses and explanatory variables used in the study.**

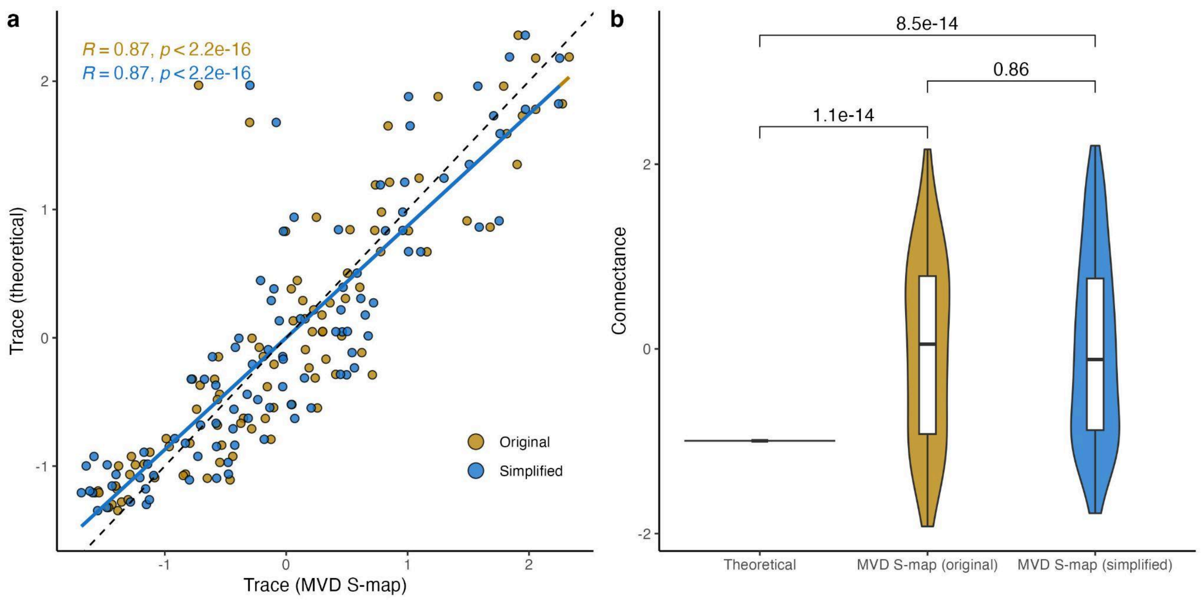


Fig. S4. **Methodological approach sensitivity analysis (MVD S-map)**. Comparing trace and connectance of dynamics created by the Multispecies-Ricker Model following Chang et al. 2021 between theoretical expectation (specified in the model) and based on coefficients estimated by the MVD S-map.

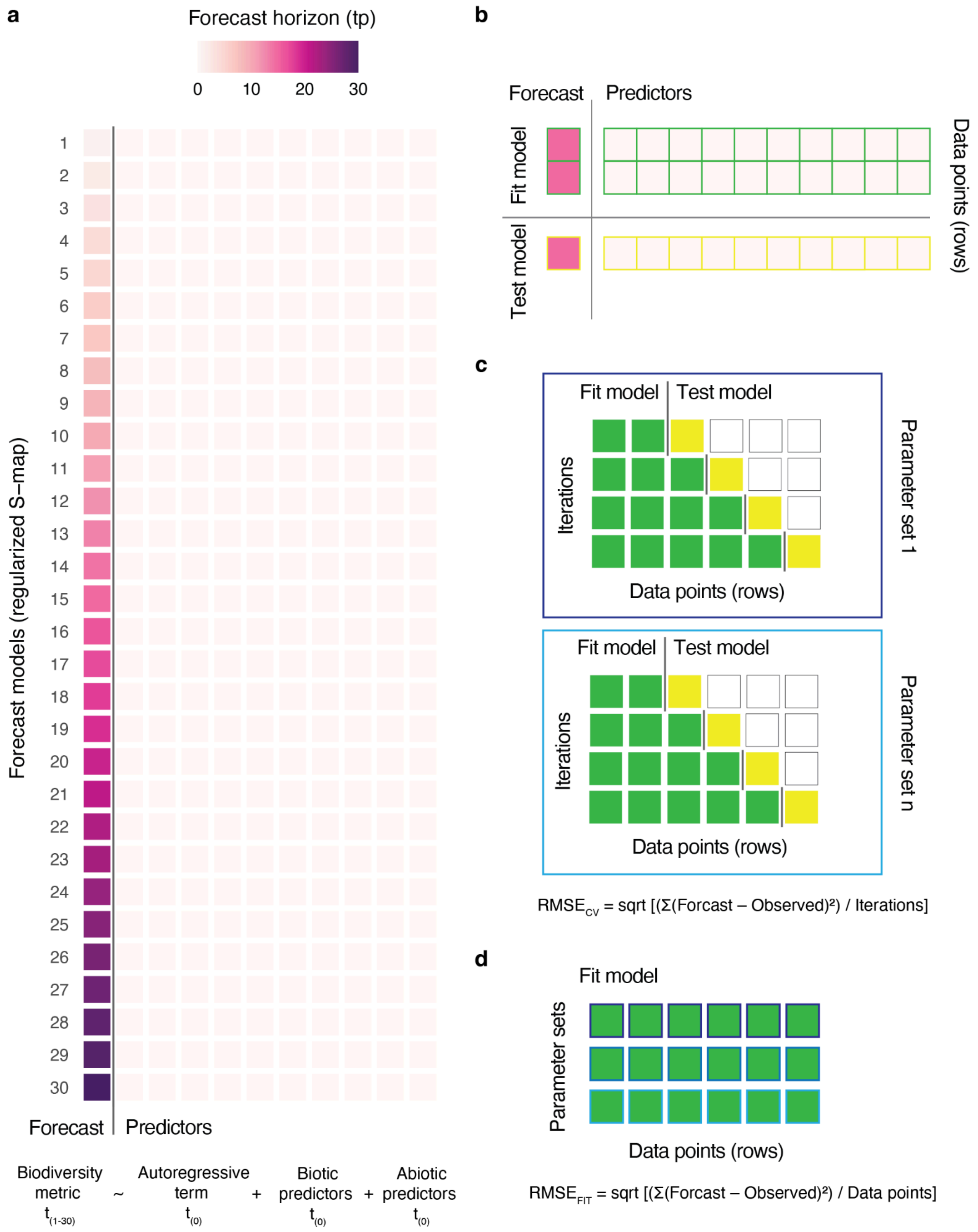


Fig. S5: **Illustration of forecasting approach.** [a] Model structure. Each forecast horizon (1-30 days) has its own model, where we predict the biodiversity metric at $t_{(1\text{-}30)}$ based on the autoregressive term, biotic predictors and abiotic predictors at $t_{(0)}$. Each square here represents a variable (forecast and the predictors).

[b] Day-forward-chaining cross-validation. The initial library to fit a model contains the first 10% data points, which are used to predict the next point to test the model. Each row represents a data point, whereas a square represents a variable. [c] Parametrization. We use different combinations of theta (model's degree of nonlinearity), lambda (strength of coefficient penalization) and alpha (regularization technique) to find the best-performing models. Model performance is evaluated using day-forward-chaining cross-validation. Predicted data points are iteratively added to the library to predict the next point, increasing the library size until all data has been used. Green squares represent data points used for model fitting, and yellow for model testing. [d] Fit model. We choose the top 5% parameter combinations (n = 6 models) based on the lowest forecast error and fit the model with the whole data set to forecast biodiversity change and average model coefficients.

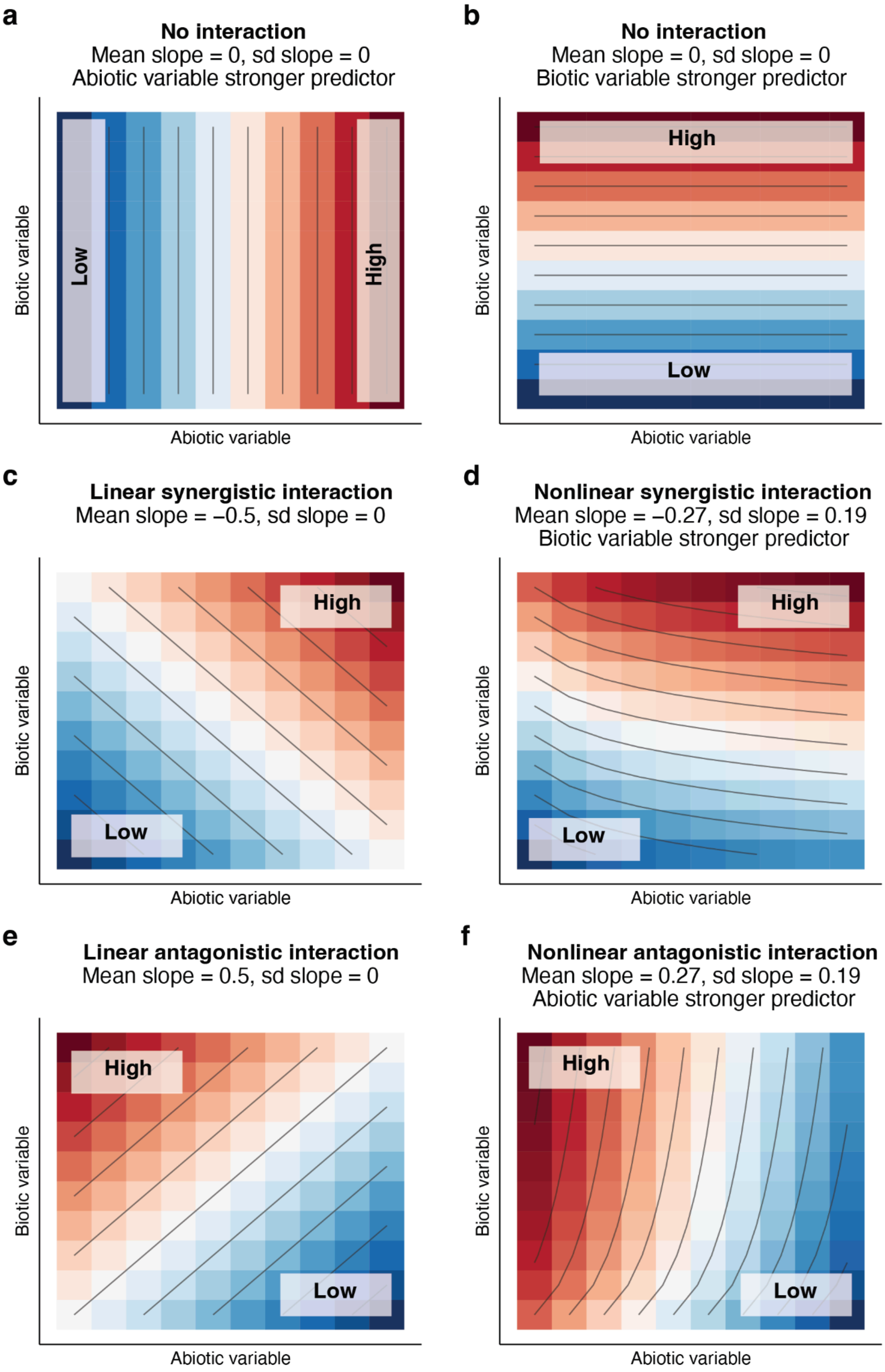


Fig. S6. **Interpreting the heatmap plots in Fig. 4 to understand the interaction between explanatory variables (six common cases).** Color code shows the predicted response (e.g., biodiversity metrics) over a gradient of biotic and abiotic explanatory variables (e.g., network connectance and temperature). The slopes indicate the relationship between the explanatory variables and their effect on the response.

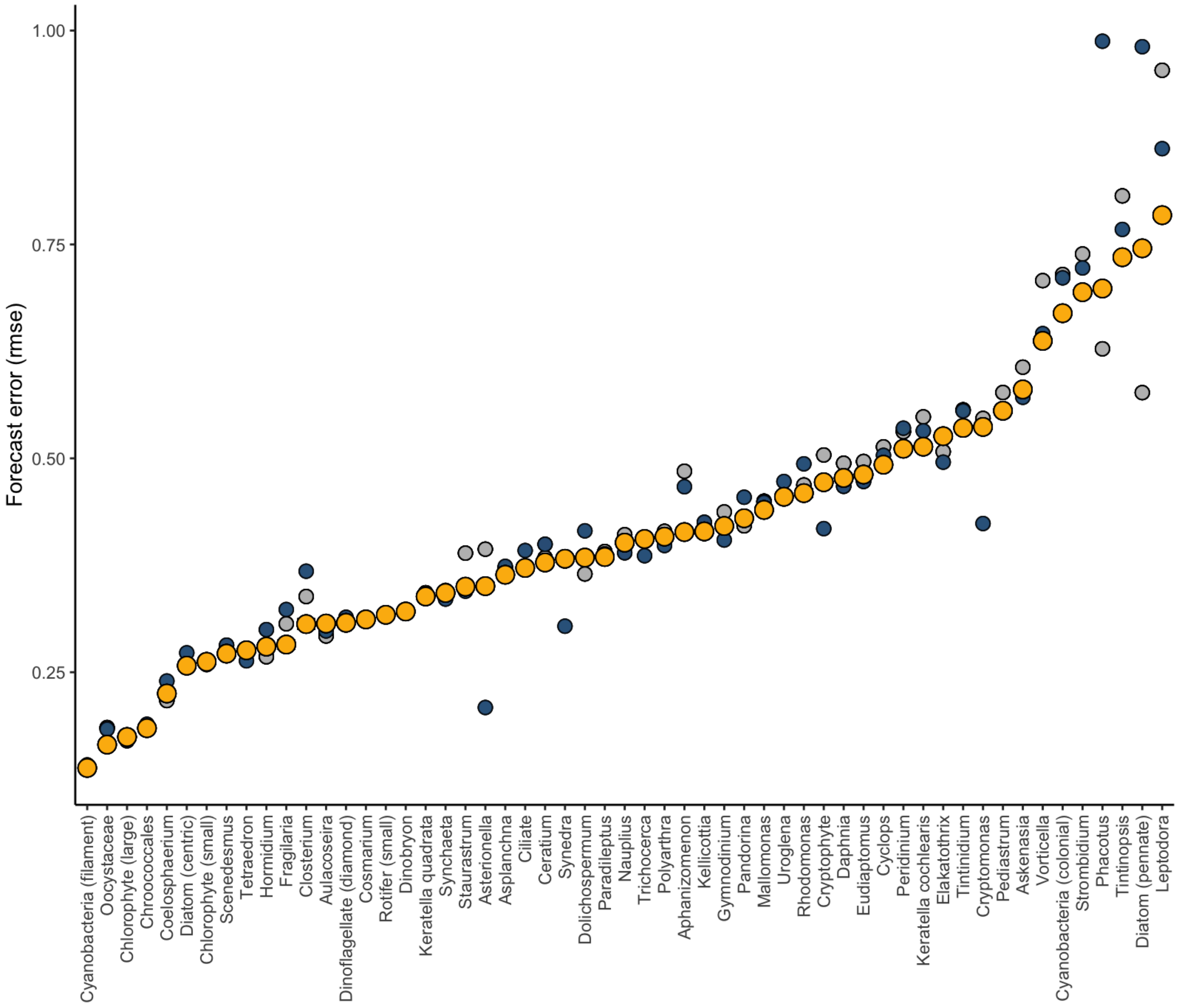


Fig. S7: **Model performance for estimating taxa interactions.** Yellow is the forecast error (rmse) for the full MVD S-map, blue is the forecast error during the parametrization and grey is the forecast error when using a constant predictor (defined by the abundance at t = 0 - abundance at t = 1).

### Tables

Tab. S1. **Summary statistics of taxonomic groups used for the study.**

| Taxon_ID | Name | Occurrence | Average abundance (ROI/s) | Size ($mm^2$) | Training images | Magnification |
|---|---|---|---|---|---|---|
| **leptodora** | Leptodora | 0.5912 | 0.0025 | 0.3251 | 276 | 0p5x |
| **asplanchna** | Asplanchna | 0.7829 | 0.0104 | 0.1263 | 679 | 0p5x |
| **cyclops** | Cyclops | 0.9638 | 0.0877 | 0.1842 | 1998 | 0p5x |
| **daphnia** | Daphnia | 0.9624 | 0.0427 | 0.341 | 1973 | 0p5x |
| **eudiaptomus** | Eudiaptomus | 0.9638 | 0.0338 | 0.2877 | 1539 | 0p5x |
| **nauplius** | Nauplius | 0.9638 | 0.1479 | 0.0546 | 2602 | 0p5x |
| **kellicottia** | Kellicottia | 0.9247 | 0.0229 | 0.0392 | 519 | 0p5x |
| **keratella_cochlearis** | Keratella cochlearis | 0.9515 | 0.0121 | 0.0185 | 782 | 0p5x |
| **keratella_quadrata** | Keratella quadrata | 0.9182 | 0.0112 | 0.0257 | 872 | 0p5x |
| **polyarthra** | Polyarthra | 0.8119 | 0.0071 | 0.026 | 127 | 0p5x |
| **rotifer_small** | Rotifer (small) | 0.9638 | 0.1737 | 0.0207 | 1107 | 0p5x |
| **synchaeta** | Synchaeta | 0.7192 | 0.0063 | 0.0102 | 371 | 0p5x |
| **trichocerca** | Trichocerca | 0.8198 | 0.005 | 0.0033 | 576 | 0p5x |
| **askenasia** | Askenasia | 0.5926 | 0.0004 | 0.0014 | 405 | 5p0x |
| **ciliate** | Ciliate | 0.9616 | 0.0039 | 0.0018 | 2078 | 5p0x |
| **paradileptus** | Paradileptus | 0.6346 | 0.0083 | 0.0292 | 581 | 0p5x |
| **strombidium** | Strombidium | 0.8683 | 0.0005 | 0.0019 | 583 | 5p0x |
| **tintinidium** | Tintinidium | 0.7627 | 0.0006 | 0.0024 | 115 | 5p0x |
| **tintinopsis** | Tintinopsis | 0.6628 | 0.0003 | 0.0015 | 51 | 5p0x |
| **vorticella** | Vorticella | 0.7012 | 0.0004 | 0.0019 | 291 | 5p0x |
| **ceratium** | Ceratium | 0.7438 | 0.0121 | 0.0231 | 1031 | 0p5x |
| **dinoflagellate_diamond** | Dinoflagellate (diamond) | 0.7779 | 0.001 | 0.0012 | 378 | 5p0x |
| **gymnodinium** | Gymnodinium | 0.9399 | 0.0022 | 0.0004 | 308 | 5p0x |
| **peridinium** | Peridinium | 0.7438 | 0.0006 | 0.0018 | 605 | 5p0x |
| **aphanizomenon** | Aphanizomenon | 0.7402 | 0.0046 | 0.0318 | 322 | 0p5x |
| **chroococcales** | Chroococcales | 0.7952 | 0.002 | 0.0005 | 183 | 5p0x |
| **coelosphaerium** | Coelosphaerium | 0.5962 | 0.0009 | 0.0016 | 450 | 5p0x |
| **cyanobacteria_colony** | Cyanobacteria (filament) | 0.9638 | 0.0065 | 0.0013 | 2066 | 5p0x |
| **cyanobacteria_filament** | Cyanobacteria (colonial) | 0.5376 | 0.0007 | 0.0021 | 551 | 5p0x |
| **dolichospermum** | Dolichospermum | 0.5579 | 0.0005 | 0.0031 | 621 | 5p0x |
| **chlorophyte_large** | Chlorophyte (large) | 0.9638 | 0.0163 | 0.0004 | 1579 | 5p0x |

| | | | | | | |
|---|---|---|---|---|---|---|
| **chlorophyte_small** | Chlorophyte (small) | 0.9638 | 0.0138 | 0.0004 | 1504 | 5p0x |
| **closterium** | Closterium | 0.8806 | 0.0014 | 0.0009 | 857 | 5p0x |
| **cosmarium** | Cosmarium | 0.6556 | 0.0014 | 0.0008 | 579 | 5p0x |
| **elakatothrix** | Elakatothrix | 0.8263 | 0.0006 | 0.0007 | 391 | 5p0x |
| **hormidium** | Hormidium | 0.7619 | 0.0017 | 0.0016 | 774 | 5p0x |
| **oocystaceae** | Oocystaceae | 0.877 | 0.002 | 0.0002 | 532 | 5p0x |
| **pandorina** | Pandorina | 0.6614 | 0.0005 | 0.0011 | 329 | 5p0x |
| **pediastrum** | Pediastrum | 0.6433 | 0.0003 | 0.0012 | 281 | 5p0x |
| **phacotus** | Phacotus | 0.9175 | 0.0012 | 0.0002 | 246 | 5p0x |
| **scenedesmus** | Scenedesmus | 0.8495 | 0.0007 | 0.0009 | 336 | 5p0x |
| **staurastrum** | Staurastrum | 0.8133 | 0.0006 | 0.0017 | 735 | 5p0x |
| **tetraedron** | Tetraedron | 0.924 | 0.0009 | 0.0002 | 174 | 5p0x |
| **asterionella** | Asterionella | 0.6556 | 0.0083 | 0.0035 | 1056 | 0p5x |
| **aulacoseira** | Aulacoseira | 0.6382 | 0.0005 | 0.0019 | 252 | 5p0x |
| **diatom_centric** | Diatom (centric) | 0.9465 | 0.0016 | 0.0008 | 1049 | 5p0x |
| **diatom_pennate** | Diatom (pennate) | 0.919 | 0.0008 | 0.001 | 321 | 5p0x |
| **fragilaria** | Fragilaria | 0.8661 | 0.0129 | 0.0297 | 1309 | 0p5x |
| **synedra** | Synedra | 0.7836 | 0.0006 | 0.0008 | 331 | 5p0x |
| **dinobryon** | Dinobryon | 0.8763 | 0.0554 | 0.0025 | 3670 | 0p5x |
| **mallomonas** | Mallomonas | 0.9624 | 0.0026 | 0.0004 | 594 | 5p0x |
| **uroglena** | Uroglena | 0.7098 | 0.0112 | 0.0368 | 1953 | 0p5x |
| **cryptomonas** | Cryptomonas | 0.9638 | 0.038 | 0.0004 | 1605 | 5p0x |
| **cryptophyte** | Cryptophyte | 0.9595 | 0.0053 | 0.0004 | 449 | 5p0x |
| **rhodomonas** | Rhodomonas | 0.9638 | 0.0114 | 0.0002 | 584 | 5p0x |

Tab. S2. **Scientific classification of taxonomic groups used for the study.**

| Taxon_ID | Empire | Kingdom | Phylum | Class | Order | Family | Genus | Species | Guild |
|---|---|---|---|---|---|---|---|---|---|
| **leptodora** | Eukaryota | Animalia | Arthropoda | Branchiopoda | Diplostraca | Leptodoridae | Leptodora | sp | Invertebrate predators |
| **asplanchna** | Eukaryota | Animalia | Rotifera | Eurotatoria | Ploima | Asplanchnidae | Asplanchna | sp | Omnivores |
| **cyclops** | Eukaryota | Animalia | Arthropoda | Maxillopoda | Cyclopoida | Cyclopidae | Cyclops | sp | Omnivores |
| **daphnia** | Eukaryota | Animalia | Arthropoda | Branchiopoda | Diplostraca | Daphniidae | Daphnia | sp | Large herbivores |
| **eudiaptomus** | Eukaryota | Animalia | Arthropoda | Maxillopoda | Calanoida | Diaptomidae | Eudiaptomus | sp | Large herbivores |
| **nauplius** | Eukaryota | Animalia | Arthropoda | Maxillopoda | | | | | Nauplia |
| **kellicottia** | Eukaryota | Animalia | Rotifera | Monogononta | Ploima | Brachionidae | Kellikottia | sp | Herbivore rotifers |
| **keratella_cochlearis** | Eukaryota | Animalia | Rotifera | Monogononta | Ploima | Brachionidae | Keratella | cochlearis | Herbivore rotifers |
| **keratella_quadrata** | Eukaryota | Animalia | Rotifera | Monogononta | Ploima | Brachionidae | Keratella | quadrata | Herbivore rotifers |
| **polyarthra** | Eukaryota | Animalia | Rotifera | Monogononta | Ploima | Synchaetidae | Polyarthra | sp | Herbivore rotifers |
| **rotifer_small** | Eukaryota | Animalia | Rotifera | Rotatoria | | | | | Herbivore rotifers |
| **synchaeta** | Eukaryota | Animalia | Rotifera | Monogononta | Ploima | Synchaetidae | Synchaeta | sp | Herbivore rotifers |
| **trichocerca** | Eukaryota | Animalia | Rotifera | Eurotatoria | Ploima | Trichocercidae | Trichocerca | sp | Herbivore rotifers |
| **askenasia** | Eukaryota | Chromalveolata | Ciliophora | Litostomatea | Cyclotrichiida | Mesodiniidae | Askenasia | sp | Ciliates |
| **ciliate** | Eukaryota | Chromalveolata | Ciliophora | | | | | | Ciliates |
| **paradileptus** | Eukaryota | Chromalveolata | Ciliophora | Litostomatea | Haptorida | Tracheliidae | Paradileptus | sp | Ciliates |
| **strombidium** | Eukaryota | Chromalveolata | Ciliophora | Oligotrichea | Oligotrichida | Strombidiidae | Strombidium | sp | Ciliates |
| **tintinidium** | Eukaryota | Chromalveolata | Ciliophora | Oligotrichea | Choreotrichida | Tintinnidiidae | Tintinnidium | sp | Ciliates |
| **tintinopsis** | Eukaryota | Chromalveolata | Ciliophora | Oligotrichea | Choreotrichida | Codonellidae | Tintinnopsis | sp | Ciliates |
| **vorticella** | Eukaryota | Chromalveolata | Ciliophora | Oligohymenophorea | Sessilida | Vorticellidae | Vorticella | sp | Ciliates |
| **ceratium** | Eukaryota | Protozoa | Dinophyta | Dinophyceae | Gonyaulacales | Ceratiaceae | Ceratium | sp | Mixotrophic flagellates |
| **dinoflagellate_diamond** | Eukaryota | Protozoa | Dinophyta | Dinophyceae | | | | | Mixotrophic flagellates |
| **gymnodinium** | Eukaryota | Protozoa | Dinophyta | Dinophyceae | Gymnodiniales | Gymnodiniaceae | Gymnodinium | sp | Mixotrophic flagellates |
| **peridinium** | Eukaryota | Protozoa | Dinophyta | Dinophyceae | Peridiniales | Peridiniaceae | Peridinium | sp | Mixotrophic flagellates |
| **aphanizomenon** | Prokaryota | Bacteria | Cyanobacteria | Cyanophyceae | Nostocales | Aphanizomenonaceae | Aphanizomenon | sp | Cyanobacteria |
| **chroococcales** | Prokaryota | Cyanobacteria | Cyanophyceae | Chroococcales | | | | | Cyanobacteria |
| **coelosphaerium** | Prokaryota | Bacteria | Cyanobacteria | Cyanophyceae | Synechococcales | Coelosphaeriaceae | Coelosphaerium | sp | Cyanobacteria |
| **cyanobacteria_colony** | Prokaryota | Bacteria | Cyanobacteria | Cyanophyceae | | | | | Cyanobacteria |
| **cyanobacteria_filament** | Prokaryota | Bacteria | Cyanobacteria | Cyanophyceae | | | | | Cyanobacteria |
| **dolichospermum** | Prokaryota | Bacteria | Cyanobacteria | Cyanophyceae | Nostocales | Aphanizomenonaceae | Dolichospermum | sp | Cyanobacteria |
| **chlorophyte_large** | Eukaryota | Plantae | Chlorophyta | | | | | | Green algae |

| | | | | | | | | | |
|---|---|---|---|---|---|---|---|---|---|
| **chlorophyte_small** | Eukaryota | Plantae | Chlorophyta | | | | | | Green algae |
| **closterium** | Eukaryota | Plantae | Charophyta | Conjugatophyceae | Desmidiales | Closteriaceae | Closterium | sp | Green algae |
| **cosmarium** | Eukaryota | Plantae | Charophyta | Conjugatophyceae | Desmidiales | Desmidiaceae | Cosmarium | sp | Green algae |
| **elakatothrix** | Eukaryota | Plantae | Charophyta | Klebsormidiophyceae | Klebsormidiales | Elakatotrichaceae | Elakatothrix | sp | Green algae |
| **hormidium** | Eukaryota | Plantae | Chlorophyta | Trebouxiophyceae | Prasiolales | Prasiolaceae | Hormidium | sp | Green algae |
| **oocystaceae** | Eukaryota | Plantae | Chlorophyta | Trebouxiophyceae | Chlorellales | Oocystaceae | | | Green algae |
| **pandorina** | Eukaryota | Plantae | Chlorophyta | Chlorophyceae | Volvocales | Volvocaceae | Pandorina | sp | Green algae |
| **pediastrum** | Eukaryota | Plantae | Chlorophyta | Chlorophyceae | Sphaeropleales | Hydrodictyaceae | Pediastrum | sp | Green algae |
| **phacotus** | Eukaryota | Plantae | Chlorophyta | Chlorophyceae | Volvocales | Phacotaceae | Phacotus | sp | Green algae |
| **scenedesmus** | Eukaryota | Plantae | Chlorophyta | Chlorophyceae | Sphaeropleales | Scenedesmaceae | Scenedesmus | sp | Green algae |
| **staurastrum** | Eukaryota | Plantae | Charophyta | Conjugatophyceae | Desmidiales | Desmidiaceae | Staurastrum | sp | Green algae |
| **tetraedron** | Eukaryota | Plantae | Chlorophyta | Chlorophyceae | Sphaeropleales | Hydrodictyaceae | Tetraedron | sp | Green algae |
| **asterionella** | Eukaryota | Chromista | Bacillariophyta | Fragilariophyceae | Tabellariales | Tabellariaceae | Asterionella | sp | Diatoms |
| **aulacoseira** | Eukaryota | Chromista | Bacillariophyta | Coscinodiscophyceae | Aulacoseirales | Aulacoseiraceae | Aulacoseira | sp | Diatoms |
| **diatom_centric** | Eukaryota | Chromista | Bacillariophyta | Bacillariophyceae | Centrales | | | | Diatoms |
| **diatom_pennate** | Eukaryota | Chromista | Bacillariophyta | Bacillariophyceae | Pennales | | | | Diatoms |
| **fragilaria** | Eukaryota | Chromista | Bacillariophyta | Bacillariophyceae | Fragilariales | Fragilariaceae | Fragilaria | sp | Diatoms |
| **synedra** | Eukaryota | Chromista | Bacillariophyta | Bacillariophyceae | Fragilariales | Fragilariaceae | Synedra | cyclopum | Diatoms |
| **dinobryon** | Eukaryota | Chromista | Ochrophyta | Chrysophyceae | Chromulinales | Dinobryaceae | Dinobryon | sp | Gold algae |
| **mallomonas** | Eukaryota | Chromista | Ochrophyta | Synurophyceae | Synurales | Mallomonadaceae | Mallomonas | sp | Gold algae |
| **uroglena** | Eukaryota | Chromista | Ochrophyta | Chrysophyceae | Chromulinales | Chromulinaceae | Uroglena | sp | Gold algae |
| **cryptomonas** | Eukaryota | Chromista | Cryptophyta | Cryptophyceae | Cryptomonadales | Cryptomonadaceae | Cryptomonas | sp | Cryptophytes |
| **cryptophyte** | Eukaryota | Chromista | Cryptophyta | Cryptophyceae | | | | | Cryptophytes |
| **rhodomonas** | Eukaryota | Chromista | Cryptophyta | Cryptophyceae | Pyrenomonadales | Pyrenomonadaceae | Rhodomonas | sp | Cryptophytes |
